# Alternative management strategy influences litter microbiome dynamics in a commercial broiler rearing system

**DOI:** 10.64898/2026.07.01.735883

**Authors:** Brett Hale, Carson Priddle, Gaurav Gajurel, Kipa Tamrakar, Makenly Coles, Luana Mendonca Dias, Peter M. Rubinelli, Elena G. Olson, Carson Arnold, Danielle Graham, Robert C. Shields, Steven C. Ricke

## Abstract

**Background:** Pre-harvest litter management is a key determinant of broiler production conditions, influencing NH₃ generation, nutrient retention, and microbial reservoirs that accumulate across production cycles. Conventional chemical and physical management strategies can support flock health, but their effects on bacterial populations are often transient and may not account for the microbial interactions that govern persistence, exclusion, and community succession. Here, we evaluated an alternative litter management strategy combining a postbiotic intervention with reduced-rate NaHSO₄ across two broiler growouts, with litter sampled at the end of each flock to determine effects on prokaryotic microbiome structure, physicochemistry, and *Enterococcus* abundance.

**Results:** Alternative management influenced observed richness and phylogenetic diversity and was associated with lower relative abundance of *Enterococcus*, including *E. cecorum* and *E. hirae* group. Sequencing-and dPCR-derived *Enterococcus* abundance estimates were correlated across samples; however, treatment-associated differences in relative abundance were not consistently accompanied by lower absolute copy number at terminal sampling, indicating changes in proportional representation rather than uniform reductions in *Enterococcus* abundance. In complementary *in vitro* assays, isolate responses to the postbiotic were strain-, concentration-, and time-dependent and were not uniformly sustained. Responses near the nominal application-mixture range were modest, whereas the largest and most persistent reductions in viable recovery occurred only at higher concentrations.

**Conclusions:** These findings demonstrate that alternative litter management can alter prokaryotic community structure and the proportional representation of selected bacterial taxa under commercial production conditions while maintaining comparable flock performance. Collectively, the field and *in vitro* observations are more consistent with broader management-associated restructuring of the litter microbial community than with sustained direct antimicrobial effects, providing a basis for microbiome-informed approaches to understanding and managing litter ecology across successive production cycles.

## Background

Broiler production is a major component of global food systems, supplying the second most consumed meat worldwide [1]. Annual output exceeds 104 million metric tons and is expected to approach 110 million metric tons in the coming years [2]. Throughout the production cycle (i.e., growout or flock), birds are reared on bedding that progressively accumulates excreta, feathers, spilled feed, moisture, and microorganisms, forming what is collectively referred to as litter. Such accumulation occurs at a scale of nearly 13 million metric tons annually across U.S. broiler operations [3]. As poultry litter ages, microbial and physicochemical transformations influence ammonia (NH₃) generation, nutrient cycling, pathogen persistence, and, consequently, flock health [4–6]. Moreover, spent litter is commonly repurposed as an organic fertilizer following flock removal, extending the impacts of management beyond the poultry house and into surrounding agricultural and environmental systems [7]. Litter thus occupies a central role in broiler production, functioning both as the immediate rearing environment and as a polymicrobial matrix responsive to management intervention.

A primary objective of broiler litter management is mitigating NH₃ volatilization, which results from microbial mineralization of nitrogenous compounds in excreta and adversely affects avian respiratory health while contributing to atmospheric emissions [6, 8]. Equally important is the pathogen load harbored within litter, which can influence early gastrointestinal colonization of chicks, contribute to disease pressure during growout, and increase downstream contamination risk during processing [9–10]. This load encompasses a broad range of clinically and agriculturally relevant taxa, including enteric pathogens such as *Salmonella* spp. and *Enterococcus* spp. and cutaneous or mucosal opportunists such as *Staphylococcus* spp., some of which possess zoonotic potential [11]. Beyond virulence, many of these organisms serve as reservoirs for antimicrobial resistance genes that accumulate within litter and may disseminate into surrounding environments following land application, with *Enterococcus* and *Staphylococcus* among the most frequently implicated genera [12–14].

Acidification represents one of the most widely adopted strategies for pre-harvest litter management, promoting the retention of ammoniacal nitrogen as nonvolatile ammonium and thereby limiting NH₃ volatilization. The benefits of this approach for bird welfare and environmental emissions are well evidenced; however, the extent to which acidification provides sustained suppression of pathogenic bacterial populations remains less certain [4, 8]. Sodium bisulfate (NaHSO₄) has been shown to transiently reduce litter pH and *Escherichia coli* abundance, with diminished impacts during growout as excreta and organic matter accumulate [15]. Other studies suggest that reduced pH alone is inadequate to suppress enteric pathogens. For example, NaHSO₄ application increased *Escherichia* abundance in top-dressed litter under commercial conditions [16], while experimentally acidified litter supported greater *Salmonella* persistence despite substantial reductions in pH [17]. These findings imply that physicochemical modification of litter may be insufficient for sustained control of pathogen-associated taxa when implemented in isolation. Furthermore, bacterial populations are influenced by microbial interactions within litter, including competitive exclusion and successional processes that shape community assembly over time [18–19]. Accordingly, complementary interventions informed by poultry litter microbial ecology may help shape microbiome structure while preserving the benefits of conventional acidification practices.

The present study evaluated an alternative litter management strategy combining a reduced NaHSO₄ regimen with a postbiotic intervention across two growouts to determine its effects on the litter prokaryotic microbiome and physicochemistry under commercial broiler rearing conditions. Prior bench-scale work showed that postbiotic-associated responses were selective and dependent on concentration and sampling time rather than broadly or consistently suppressive, with the most notable differences occurring among KF Streptococcus agar- and Baird-Parker agar-recoverable populations representing presumptive enterococci and staphylococci, respectively [20]. Similar patterns were observed under commercial conditions, where application of the postbiotic in conjunction with reduced NaHSO₄ was associated with shifts in microbiome composition and lower *Staphylococcus* relative abundance following flock removal [21]. Here, full-length 16S rRNA gene sequencing was used to characterize management-associated differences in the litter microbiome at the conclusion of each growout. Patterns in *Enterococcus* abundance were subsequently evaluated using genus- and species-level digital PCR (dPCR) assays and complemented with *in vitro* testing using reference and poultry-derived isolates. The litter physicochemical profile was additionally characterized and used to contextualize relationships among management, microbial community structure, and taxon abundance. Observations herein advance our understanding of how alternative litter management influences poultry litter ecology and support the development of sustainable broiler production practices.

## Materials and Methods

### Animal ethics statement

Flocks remained under existing integrator management throughout the trial, and no experimental manipulation of animals, animal handling, experimental infection, euthanasia, or collection of animal tissues was performed. Therefore, institutional animal ethics approval was not required. All litter sampling was performed in accordance with the biosecurity and personal protective equipment requirements of the poultry integrator.

### Experimental design

This study was conducted at a commercial broiler operation in DeQueen, Arkansas, USA and spanned three consecutive growouts. Six mechanically ventilated broiler houses with 167.6 × 15.2 m dimensions (∼2,555 m²) were chosen as independent experimental units based on litter age (two growouts prior to study initiation) and comparable performance history (**Supplementary Figure 1**). Bedding comprised locally sourced pine chips (*Pinus* sp.), and litter had been treated with ∼488 g m⁻² of an NaHSO₄-based amendment (Jones-Hamilton Co., Maumee, OH, USA) in prior growouts. Between production cycles, 5 to 10 cm of cake [22] was removed.

Seventy-two hours before flock placement, three houses received the postbiotic intervention (AgriGro Inc., Doniphan, MO, USA) at ∼1.42 L per 93 m², while the remaining three houses served as standard-management controls. In treated houses, NaHSO₄ was applied at half the standard rate (∼244 g m⁻²), while control houses received the full rate (∼488 g m⁻²) as described above. Ross 700 chicks (Aviagen, Huntsville, AL, USA) were placed at 26,600 birds per house in growout 1 (∼10.4 birds m⁻²) and 27,200 birds per house in growouts 2 and 3 (∼10.6 birds m⁻²). All other production parameters were held constant across treatments throughout the growouts.

Ventilation and nutritional regimens followed those described by Olson et al. [21]. Fans were controlled on a 300-s cycle, with on-times starting at ≥ 60 s early in the growout and increasing to ∼240 s by flock removal. Brooding temperature began at ∼32 °C and was gradually reduced to 18 to 20 °C by flock removal, with relative humidity maintained at 50-70%. A standard four-phase broiler feeding program was used with starter, grower, finisher, and withdrawal diets [23].

### Sample collection

At broiler physiological maturity (approximately 59 days of age), five litter subsamples were collected to a depth of 10 cm and then combined in a vessel and thoroughly homogenized with a sanitized trowel. As spatial heterogeneity impacts both microbial [5, 24] and physicochemical [25] litter dynamics, five composite samples per house were collected spanning areas of anticipated highest bird density (**Figure 1**), and sample location (hereafter, “position”) was included as a factor in downstream statistical analyses. Each composite was split into two sterile 50-mL conical tubes and one resealable polyethylene bag. Tubes were placed on dry ice within 5 min of collection and stored at - 80 °C until DNA isolation. Bagged subsamples were held at ambient temperature until physicochemical analysis. Additionally, one composite sample per house was collected by the integrator prior to study initiation to capture baseline physicochemistry. Houses were windrowed prior to sample collection in the third growout; therefore, only samples from growouts 1 and 2 were carried forward to subsequent steps.

**Figure 1.**
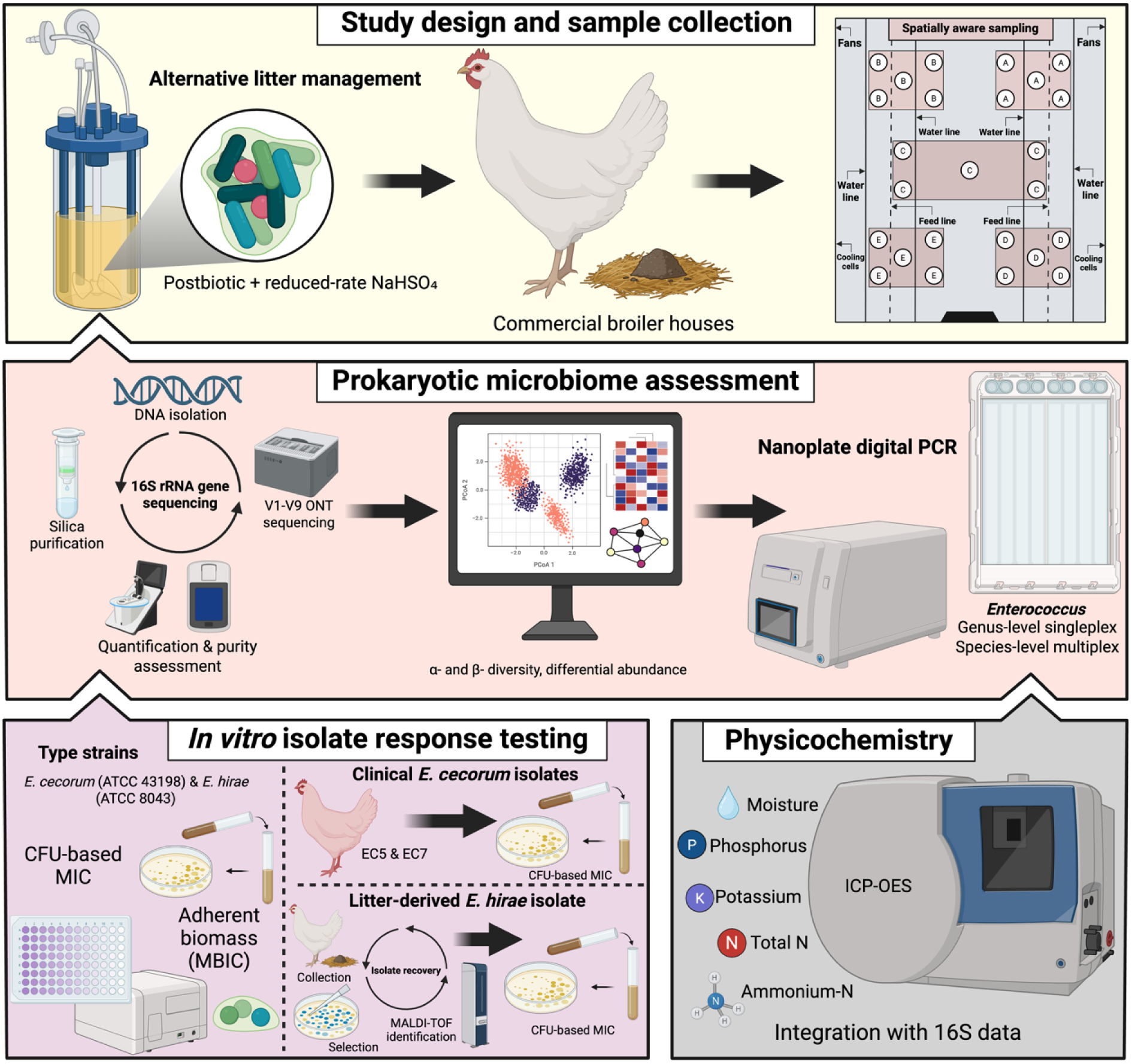
Study overview. Alternative litter management consisted of postbiotic intervention with reduced-rate NaHSO₄ application in commercial broiler houses. Litter was sampled using a spatially aware design and analyzed using full-length 16S rRNA gene sequencing, nanoplate digital PCR targeting *Enterococcus* spp., *in vitro* isolate response assays, and physicochemical profiling. This figure was created at https://BioRender.com under agreement number SA29WIM3R2.

### DNA isolation and full-length 16S rRNA gene sequencing

DNA was extracted from 250 mg of each composite sample using the DNeasy PowerSoil Pro QIAcube Kit (Qiagen, Hilden, Germany; Cat. #47126) following the manufacturer’s protocol. Mechanical homogenization was carried out with a QIAgen TissueLyser III at 25 Hz for 10 min. All downstream purification steps were automated on the QIAcube Connect system. To control for potential reagent-derived contamination, extraction-negative controls were included in triplicate as recommended by Lyte et al. [26–27].

Given the prevalence of PCR inhibitors in poultry litter matrices [28–29], samples were subjected to a secondary silica-based purification step using the DNeasy PowerClean Pro Cleanup Kit (Cat. #12997-50). The DNA concentrations were quantified using a Qubit fluorometer (Thermo Fisher Scientific, Waltham, MA, USA) with the dsDNA High-Sensitivity Assay Kit (Cat. #Q32851). DNA purity was evaluated by measuring absorbance ratios at 260/280 nm and 260/230 nm using a NanoDrop OneC spectrophotometer (Thermo Fisher Scientific).

Full-length (V1-V9) 16S rRNA gene sequencing was performed with Oxford Nanopore Technologies (ONT; Oxford, UK) chemistry. Amplicons were generated using a universal 16S primer set consisting of 27F (5′-AGRGTTYGATYMTGGCTCAG-3′) and 1492R (5′-RGYTACCTTGTTACGACTT-3′) [30]. Amplification was performed using Q5® High-Fidelity 2X Master Mix (New England Biolabs, Ipswich, MA, USA; Cat. #M0492L) with the following thermocycling conditions: initial denaturation at 98 °C for 30 s, followed by 25 cycles of 98 °C for 10 s, 56 °C for 30 s, and 72 °C for 1 min. A final extension was performed at 72 °C for 2 min. Amplicons were purified using solid-phase reversible immobilization bead cleanup (Beckman Coulter, Brea, CA, USA; Cat. #B23319).

Library preparation was carried out using the ONT SQK-NBD114 native barcoding kit following the manufacturer’s instructions. Sequencing was performed on the GridION platform using FLO-MIN114 Spot-ON flow cells (R10.4 chemistry) with a translocation speed of 400 bases per second.

### ONT basecalling and taxonomic classification

Basecalling and barcode trimming were performed directly on the GridION using MinKNOW (v24.02.16) and Dorado (v7.3.11), applying the super-accurate basecalling model. Taxonomic classification was conducted using the Epi2Me Labs 16S workflow (wf-16s; available at https://github.com/epi2me-labs/wf-16s). Reads were filtered to retain those between 800 and 2,000 bp in length and classified using minimap2 (v2.26-r1175) with a minimum percent identity of 95% and a minimum reference coverage of 90%. Classifications were performed against a curated 16S/18S reference database assembled by Epi2Me Labs, which includes NCBI RefSeq [31] sequences and is formatted for Kraken2 (v2.1.3) [32]. The workflow also incorporated the supporting software pysam (v0.21.0), pandas (v2.0.3), fastcat (v0.15.1), samtools (v1.18), and taxonkit (v0.15.1). Per-sample raw, length-filtered, classified, and analyzed read counts are provided in **Supplementary File 1**. Species-level assignments for *E. cecorum* and *E. hirae* were subsequently verified by realigning classified reads (all *E. hirae* reads and a random subsample of 200 *E. cecorum* reads) to all near-full-length (≥ 1,400 bp) *Enterococcus* type-strain 16S rRNA gene sequences in NCBI RefSeq (BioProject PRJNA33175) using the ‘pairwiseAlignment’ function in *pwalign* (v1.4.0) [33], with assignments considered resolved when percent identity to the assigned species exceeded that of the closest alternative species by more than 0.5 percentage points.

### Microbiome statistical analysis

Study metadata, taxonomy, and species-level feature counts were imported into *RStudio* (v4.2.2) [34] and used to construct a phyloseq object with the *phyloseq* R package (v1.42.0) [35]. Thereafter, we identified and removed contaminant taxa from the dataset using the ‘isContaminant’ function in the *decontam* package (v1.13) [36] based upon feature representation in negative extraction controls. Family-level sample composition was visualized thereafter with *ggplot2* (v3.5.1) [37].

To determine global patterns of taxonomy and putative traits in the dataset, we performed multiple sequence alignment (MSA) using the MUSCLE algorithm via the ‘msaMuscle’ function from the *msa* package (v1.30.1) [38], with parameters set to "upgma" (Unweighted Pair Group Method with arithmetic mean) guide-tree clustering, default gap penalties, and default substitution matrix. The resulting MSA was converted to a phyDat object using the ‘as.phyDat’ function from the *phangorn* package (v2.11.1) [39]. This phyDat object was then used to construct a distance matrix with the ‘dist.ml’ function, and a neighbor-joining tree was estimated using the ‘nj’ function from the *ape* package (v5.8) [40]. To assess clade stability, 1,000 bootstrap replicates were performed using the ‘bootstrap.phyDat’ function in *phangorn*, with each replicate generating a neighbor-joining tree from a resampled alignment. Bootstrap support values for each internal node were then calculated using the ‘prop.clades’ function. The final cladogram was visualized with the *ggtree* package (v3.12.0) [41].

Strain-level trait annotations were obtained from the BacDive database [42] with the ‘retrieve’ function of the *BacDive* R package (v0.8.0). A custom function was applied to retrieve and collate information on gram stain, cell shape, oxygen tolerance, antibiotic resistance, enzyme utilization, and metabolite utilization for all strains of each species. Pie charts for each annotation were constructed using *ggplot2*.

We next assessed treatment effects on microbiome structure and composition. Within-sample (α) diversity was estimated by subsampling to the minimum library size across 1,000 iterations using the ‘rarefy_even_depth’ *phyloseq* function. Shannon index [43], observed richness, and inverse Simpson [44] indices were calculated using the ‘estimate_richness’ function in *phyloseq*. Furthermore, Faith’s phylogenetic diversity [45] was inferred with the ‘pd’ function from the *picante* package (v1.8.2) [46]. Final α-diversity values for each sample were taken as the mean across the 1,000 iterations.

Prior to model fitting, data distributions were assessed with a Shapiro-Wilk test [47] from the *stats* package (v4.2.2), and by qualitative assessment of histograms generated with *ggplot2*. Normal, gamma, and log-normal models were then fitted using ‘fitdistr’ from the *MASS* package (v7.3-61) [48] and compared using the Akaike Information Criterion (AIC) implemented with the *stats* function ‘AIC’. Generalized linear mixed models (GLMMs) were constructed in *glmmTMB* (v1.1.9) [49], specifying treatment, growout, position, and the treatment-growout interaction as fixed effects, and including house as a random intercept. For metrics best fit by a log-normal distribution, the response was log-transformed and modeled using a GLMM with a Gaussian error distribution. Pairwise treatment comparisons within each growout were obtained from estimated marginal means leveraging the *emmeans* package (v1.10.4) [50].

Model diagnostics were generated with the *DHARMa* package (v0.5.0) [51]. We simulated 1,000 randomized-quantile residuals via ‘simulateResiduals’ and assessed dispersion (‘testDispersion’), outliers (‘testOutliers’) and overall quantile fit (‘testQuantiles’) for each GLMM. Models were deemed acceptable when dispersion did not differ significantly from unity (*p* > 0.05) and no systematic outlier or quantile deviations were detected.

Subsequently, feature counts were normalized by cumulative sum scaling (CSS) with *metagenomeSeq* (v1.46.0) [52]. Global Bray-Curtis [53], weighted UniFrac, and unweighted UniFrac [54] distance matrices were then constructed and assessed via Principal Coordinates Analysis (PCoA) with the ‘ordinate’ *phyloseq* function, followed by visualization with *ggplot2*. Compositional dissimilarity was inferred statistically by permutational multivariate analysis of variance (PERMANOVA) with collective main effects of treatment, growout, and position tested using distance ∼ Treatment + Growout + Position, and the treatment-growout interaction tested using distance ∼ Treatment * Growout + Position. The PERMANOVA was performed with the ‘adonis2’ function from the *vegan* R package (v2.6-6.1) [55] using marginal tests (by = "margin") and factor-specific restricted permutation schemes: whole-house permutations for treatment and treatment-growout effects, growout permutations within house, and position permutations within house while retaining positions across growouts. Homogeneity of group dispersions was examined using the ‘betadisper’ function from *vegan*. The statistical significance of differences in distances to group medians was assessed with both the ‘anova’ and ‘permutest’ functions from *vegan*, with the corresponding restricted permutation structure used for permutation tests. Treatment effects were also evaluated independently within the first and second growouts using distance ∼ Treatment + Position. For these analyses, treatment effects were evaluated by permuting houses as whole experimental units, whereas position effects were evaluated using position permutations within house. The PERMANOVA variance attributable to each fixed effect was visualized using the *ComplexHeatmap* package (v2.20.0) [56].

To identify differentially abundant features between treatment levels, we employed the R package *MaAsLin2* (Microbiome Multivariable Associations with Linear Models) (v1.18.0) [57]. The CSS-normalized count matrices were modeled using negative binomial (NEGBIN), zero-inflated negative binomial (ZINB), and compound poisson linear models (CPLM). All models incorporated treatment, position, and growout as fixed effects, and house as a random intercept. To increase statistical power, we also constructed genus-level phyloseq objects using the ‘tax_glom’ function in *phyloseq* and repeated the *MaAsLin2* analysis with the described parameters. Features with Benjamini-Hochberg-adjusted *q*-values ≤ 0.05 [58] were considered differentially abundant. Analyses were repeated independently within each growout (treatment and position as fixed effects; house as a random intercept), and the log-normalized false discovery rate (FDR; -sign[coefficient]*log[*q*-value]) [21] for differentially abundant features was visualized using *ComplexHeatmap*.

### dPCR quantification of *Enterococcus* spp

Nanoplate dPCR assays were developed to quantify *Enterococcus* spp., *E. cecorum*, and *E. hirae*. The genus-level assay targeted the 23S rRNA gene using primer sequences from Ryu et al. [59]. The *E. cecorum* assay targeted *sodA* (encoding superoxide dismutase) with a previously published primer-probe combination [60], leveraging a 6-carboxyfluorescein (FAM) reporter and a BHQ1 quencher. The *E. hirae* assay was adapted from Kim et al. [61] and targeted *sdf* (encoding sodium/glutamate symport protein) with a hexachloro-fluorescein (HEX) reporter and BHQ1 quencher.

Assay design, optimization, and validation were conducted in accordance with the Minimum Information for Publication of Quantitative Digital PCR Experiments (dMIQE) guidelines [62] and recommendations of de Korne-Elenbaas et al. [63]. Primer and probe specificity were evaluated *in silico* using NCBI Primer-BLAST [64]. Experimental optimization included gradient-based determination of annealing temperature and template DNA input. Analytical sensitivity and quantification performance were established by determining the limit of blank (LOB), limit of detection (LOD), and limit of quantification (LOQ) using no-template controls and synthetic double-stranded DNA fragments (gBlocks; Integrated DNA Technologies, Coralville, IA, USA) consisting of the target amplicon flanked by 75 bp on each side. Assay performance was evaluated based on fluorescence cluster separation using a peak resolution metric defined as the difference in mean fluorescence intensity between positive and negative partitions divided by the combined dispersion of the clusters [63]. For the multiplex assay, potential primer-probe interactions and secondary structures were assessed using the IDT OligoAnalyzer tool, with pairwise dimerization inferred from Gibbs free energy. Primer and probe sequences are provided in **Supplementary Table 1**.

All assays were performed on a QIAcuity One 5-plex System (Qiagen, Hilden, Germany) using QIAcuity Nanoplates (8,500 partitions per reaction; Cat. #250021). The genus-level singleplex assay used the QIAcuity EvaGreen PCR Kit (Cat. #250112) with 4 ng input DNA per reaction and the following conditions: 2 min PCR initial heat activation, 40 cycles of a two-step thermal profile of 15 s at 95°C for denaturation, and 60 s at 50°C for annealing and extension, followed by cooling for 5 min at 40°C. The 15 µL dPCR reaction for singleplex consisted of 3x EvaGreen PCR Master Mix (5 µL), 10x primer mix (at 0.4 µM), DNA template, and nuclease-free H_2_O. The multiplex assay used the QIAcuity Probe PCR Kit (Cat. #250102) with 10 ng input DNA and the following conditions: 2 min PCR initial heat activation, 40 cycles of a two-step thermal profile of 15 s at 95°C for denaturation, and 60 s at 50°C for annealing and extension. The 15 µL dPCR system for multiplex consisted of 4x Probe PCR master mix (3.5 µL), 10x primer-probe mix (primers at 0.8 µM and probes at 0.4 µM), DNA template and nuclease-free H2O. All samples were run in technical duplicate.

Absolute abundance was expressed as copies of target per g of litter based on Poisson-corrected partition counts. Treatment effects were evaluated with GLMMs as described for α-diversity analyses, with the exception that species-level data were modeled using a Tweedie distribution. Agreement between log₁₀ dPCR abundances and log₁₀ CSS-normalized ONT counts was assessed with Spearman rank correlation [65] using the *stats* functions ‘cor’ and ‘cor.test’, with *p*-values derived from the corresponding t-statistic. Data were visualized with *ggplot2*.

### *Enterococcus* reference-strain *in vitro* response assays

To characterize changes in adherent biomass following postbiotic exposure, we adapted a crystal-violet microtiter plate assay based on conventional minimum biofilm inhibitory concentration (MBIC) methods for Gram-positive bacteria [66–67]. Briefly, the postbiotic (50% stock solution prepared in phosphate-buffered saline [PBS]) was serially two-fold diluted in tryptic soy broth (TSB) supplemented with 0.025% glucose to obtain final concentrations ranging from 24% to 0.75% (v/v). Wells containing only culture medium served as the negative control, whereas wells containing bacterial inoculum without treatment served as the untreated positive control (BCT). Vancomycin (8 µg/mL) was included as a reference antimicrobial control. Additionally, wells containing each postbiotic concentration without bacterial inoculum were included as blanks for background subtraction. Type strains of *E. hirae* (ATCC 8043) and *E. cecorum* (ATCC 43198) were obtained from the American Type Culture Collection (ATCC; Manassas, VA, USA) and propagated according to the supplier’s recommendations. Overnight cultures prepared in TSB supplemented with 0.025% glucose were adjusted to 1 × 10⁸ colony-forming units (CFU)/mL prior to inoculation.

For each well, 100 µL of the standardized bacterial inoculum was mixed with 100 µL of the corresponding postbiotic dilution, resulting in a final volume of 200 µL per well. Plates were incubated aerobically at 37°C for 24 h to allow adherent biomass formation. Thereafter, planktonic cells were carefully removed and washed with Milli-Q water to eliminate non-adherent cells. Adherent biomass was stained with 200 µL of 0.005% crystal violet solution for 15 min at room temperature. Excess stain was removed by washing with Milli-Q water, and the bound crystal violet was solubilized with 200 µL of 7% acetic acid [68]. Biomass was quantified by measuring absorbance at 570 nm using a Synergy H1 microplate reader (BioTek Instruments, Winooski, VT, USA). Blank-corrected absorbance values were normalized to the untreated control, which was standardized to represent 100% adherent biomass. All experiments were performed in quadruplicate and repeated independently on three separate occasions (*n* = 12/group).

Treatment effects were analyzed independently for each strain using one-way ANOVA implemented with the ‘aov’ function from the *stats* package, with treatment specified as the fixed effect. When a significant treatment effect was detected (*p* < 0.05), pairwise comparisons were conducted using Tukey’s honestly significant difference (HSD) test via the ‘TukeyHSD’ function from *stats*. Compact letter displays were generated using the ‘multcompLetters4’ function from the *multcompView* package (v0.1-11) [69]. Data were visualized as percent reduction in adherent biomass relative to the untreated control within each strain and plotted using *ggplot2*.

Furthermore, CFU-based minimum inhibitory concentration (MIC) assays were conducted for *E. hirae* and *E. cecorum* using the same treatment structure, controls, and serial dilution scheme described above, with modifications for longitudinal viable-cell enumeration. Overnight cultures were prepared by growing *E. hirae* in brain heart infusion (BHI) broth and *E. cecorum* in BHI supplemented with 1% yeast extract. Prior to inoculation, bacterial cultures were normalized to an optical density at 600 nm (OD600) of 0.5 and inoculated at an approximate final concentration of 1 × 10⁶ CFU/mL.

Bacterial viability was monitored from day 0 through day 4. At each sampling time point, 10 µL aliquots were collected from each reaction mixture, serially diluted ten-fold, and spot plated onto BHI agar for viable colony enumeration. Plates inoculated with *E. cecorum* were incubated overnight under CO₂-enriched conditions, whereas *E. hirae* plates were incubated overnight aerobically at 37°C. Colony-forming units were enumerated following incubation. Data visualization and statistical analyses were performed as described for MBIC assays

### *Enterococcus* clinical isolate *in vitro* response assays

We evaluated two additional *E. cecorum* isolates at the University of Arkansas (John K. Skeeles Poultry Health Laboratory, Fayetteville, AR), designated EC5 and EC7. Both isolates were recovered from broiler chickens exhibiting clinical symptoms within the same geographic region in which the present study was conducted [70]. EC5 has been associated with reduced body weight gain, systemic colonization, lesion development (i.e., pericarditis, hydropericardium, splenomegaly, and femoral head osteomyelitis), and altered cecal microbiome composition. In contrast, EC7 has been associated with more severe systemic infection, increased recovery from internal organs, and pronounced cardiac pathology including pericarditis, focal heart necrosis, and hydropericardium [70].

For *in vitro* assays, EC5 and EC7 were incubated statically at 37°C under microaerophilic conditions for 24 h, then diluted to a target inoculum of 1 × 10⁴ CFU/mL. Each isolate was inoculated into TSB containing the postbiotic at five concentrations (1.5%, 3%, 6%, 12%, and 24% v/v), alongside untreated isolate-plus-TSB and TSB-only controls. Five biological replicates were prepared per treatment per strain (*n* = 5). Cultures were incubated as described above using Campy environment sachets in sealed chambers or a 5% CO₂ incubator. Viable cell counts were determined at 6, 12, and 24 h post-inoculation by serial ten-fold dilution in sterile saline followed by drop plating onto CHROMagar Orientation agar. Plates were dried and incubated prior to colony enumeration, and viable counts were expressed as log₁₀ CFU/mL.

An *E. hirae* isolate was obtained using a recently described poultry litter microcosm model [20] at the University of Wisconsin-Madison (Meat Science and Animal Biologics Discovery facility, Madison, WI). Litter samples inoculated with poultry-derived cecal microbiota were serially diluted in anaerobic diluting solution and enumerated via spot plating on Kenner Fecal Streptococcus Agar. Plates were incubated under microaerophilic conditions at 37-42°C, and colonies consistent with *Enterococcus* morphology were submitted to the Wisconsin Veterinary Diagnostic Laboratory for species-level identification using matrix-assisted laser desorption/ionization time-of-flight (MALDI-TOF) mass spectrometry. The recovered *E. hirae* isolate was subsequently evaluated in modified CFU-based MIC assays using the postbiotic.

For *in vitro* testing, overnight cultures of the *E. hirae* isolate were grown microaerobically in TSB and quantified using a Petroff-Hausser counting chamber prior to dilution to 1 × 10⁷ CFU/mL. Aliquots containing 3 × 10⁴ cells were inoculated into treatment wells containing the postbiotic concentrations described for the *E. cecorum* clinical isolates. Microplates were incubated at 37°C under microaerophilic conditions, and viable counts were quantified at the same sampling intervals described previously. At each sampling time point, cultures were serially diluted ten-fold in sterile diluent and 10 µL aliquots of each dilution were spot plated onto CHROMagar Orientation agar. Agar plates were incubated for 24 h at 37°C under microaerophilic conditions prior to colony enumeration. Data analysis and visualization were performed as described for type strains.

### Physicochemical analysis

Ammonium-N (NH₄⁺), total N, phosphorus (P), potassium (K), and moisture were quantified following the Recommended Methods of Manure Analysis (RMMA) [71] and in accordance with US EPA SW-846. Following sample homogenization, moisture was determined gravimetrically by oven drying at 105 °C to constant mass and reported as percent wet basis (% wb). NH₄⁺ was quantified on 2 M KCl extracts using a flow-injection autoanalyzer with salicylate colorimetry. Total N was measured after sample combustion per RMMA recommendations. P and K were determined after nitric acid digestion, with concentrations determined by inductively coupled plasma-optical emission spectrometry. Final elemental measurements were converted to kg metric ton⁻¹ and expressed on a dry-matter basis.

Data visualization and statistical analyses were performed as described for α diversity, with the exception that baseline measurements (one composite sample per house) were compared between treatments using exact two-sided Mann-Whitney U tests [72] implemented with the *stats* function ‘wilcox.test’. To assess the overall degree of correlation between microbial community structure and the physicochemical profile, we applied a Procrustean randomization test (PROTEST) [73] using the vegan ‘protest’ function with 9,999 permutations. Principal coordinate configurations were derived for each community distance (Bray-Curtis, weighted UniFrac, unweighted UniFrac) and compared to a configuration from the Euclidean distance of the z-standardized environmental matrix. We further identified the physicochemical variables most strongly associated with each distance metric using Mantel tests [74] implemented with the vegan ‘mantel’ function (Spearman’s rank correlation; 9,999 permutations), testing each standardized environmental variable independently against each community distance. For both analyses, permutations were restricted within house while retaining recurring positions across growouts.

To test how the environment was associated with β-diversity after adjusting for primary experimental factors, we ran PERMANOVA as described previously, with treatment, growout, position, and the treatment-growout interaction included in the model and physicochemical properties added as covariates (9,999 restricted permutations). The linear influence of the physicochemical profile on overall community structure was then investigated using partial distance-based redundancy analysis (db-RDA) [75] via the ‘dbrda’ function in the *vegan* package, with each physicochemical property tested independently as the constraint variable. To account for experimental structure, house, growout, treatment, and position were included in the model as conditioning variables. Each physicochemical variable was tested by permutation ANOVA using the same restricted permutation structure, and adjusted R² was used to quantify the variation attributable to the environmental constraint. The Mantel, PERMANOVA, and db-RDA outputs were visualized with *ggplot2*.

Taxon-environment associations were evaluated with *MaAsLin2* as described for differential abundance analysis, modeling all physicochemical variables and experimental covariates as fixed effects and house as a random intercept. Significant associations were displayed with *ComplexHeatmap*.

## Results

### Trial design and flock performance

The objective of this study was to determine impacts of alternative litter management on the prokaryotic microbiome in a commercial broiler operation. To do so, we compared three houses with postbiotic + half-rate NaHSO₄ with three houses receiving full-rate NaHSO₄ (common litter treatment regimen) across three consecutive nine-week growouts. Metrics assessed included per-house mortality, litter physicochemistry, *Enterococcus* abundance, and litter microbiome dynamics on the last day of two consecutive flocks. Cumulative mortality for the two growouts preceding study initiation indicated comparable performance across groups (*p* = 0.83; designated Control houses = 2,233 ± 710; designated Alternative houses = 2,338 ± 906 [mean ± 1 SD]) (**Supplementary Figure 1a**), providing an appropriate baseline for detecting treatment effects. During the study, cumulative mortality did not differ statistically between treatments yet was numerically lower in alternatively managed houses compared with controls (*p* = 0.28; Control = 2,105 ± 320; Alternative = 1,986 ± 362) (**Supplementary Figure 1b**-**d**).

### Full-length 16S sequencing captured taxonomic and trait diversity in broiler litter

Composite litter samples were collected on the day preceding flock removal for growouts 1 and 2 and subjected to V1-V9 16S rRNA gene sequencing via ONT chemistry. A collective 1,608,028 full-length reads were retained after filtering (1,486 ± 165 bp), corresponding to 26,800 ± 7,202 reads per sample (**Supplementary File 1**; **Supplementary Figure 2a**). From these, we obtained 1,573 features (272 ± 61 per sample) spanning 13 phyla, 31 classes, 71 orders, 161 families, and 466 genera (**Supplementary Figure 2b**). At phylum level, the litter microbiome was dominated by Firmicutes (Bacillota) and Actinobacteria (Actinomycetota), with lower ranks being most represented by Bacilli (class), Lactobacillales (order), Carnobacteriaceae (family), and *Atopostipes* (genus) (**Supplementary Figure 2c**).

The feature sequences were aligned and used to construct a neighbor-joining tree to infer phylogeny-aware microbiome metrics, demonstrating clades consistent with taxonomic assignments (**Figure 2a**). The BacDive database was then used to acquire species metadata, wherein 74.6% were Gram-positive, 11.1% Gram-negative, 1.0% Gram-variable (inconsistent staining due to cell wall characteristics), and 13.3% not reported.

**Figure 2.**
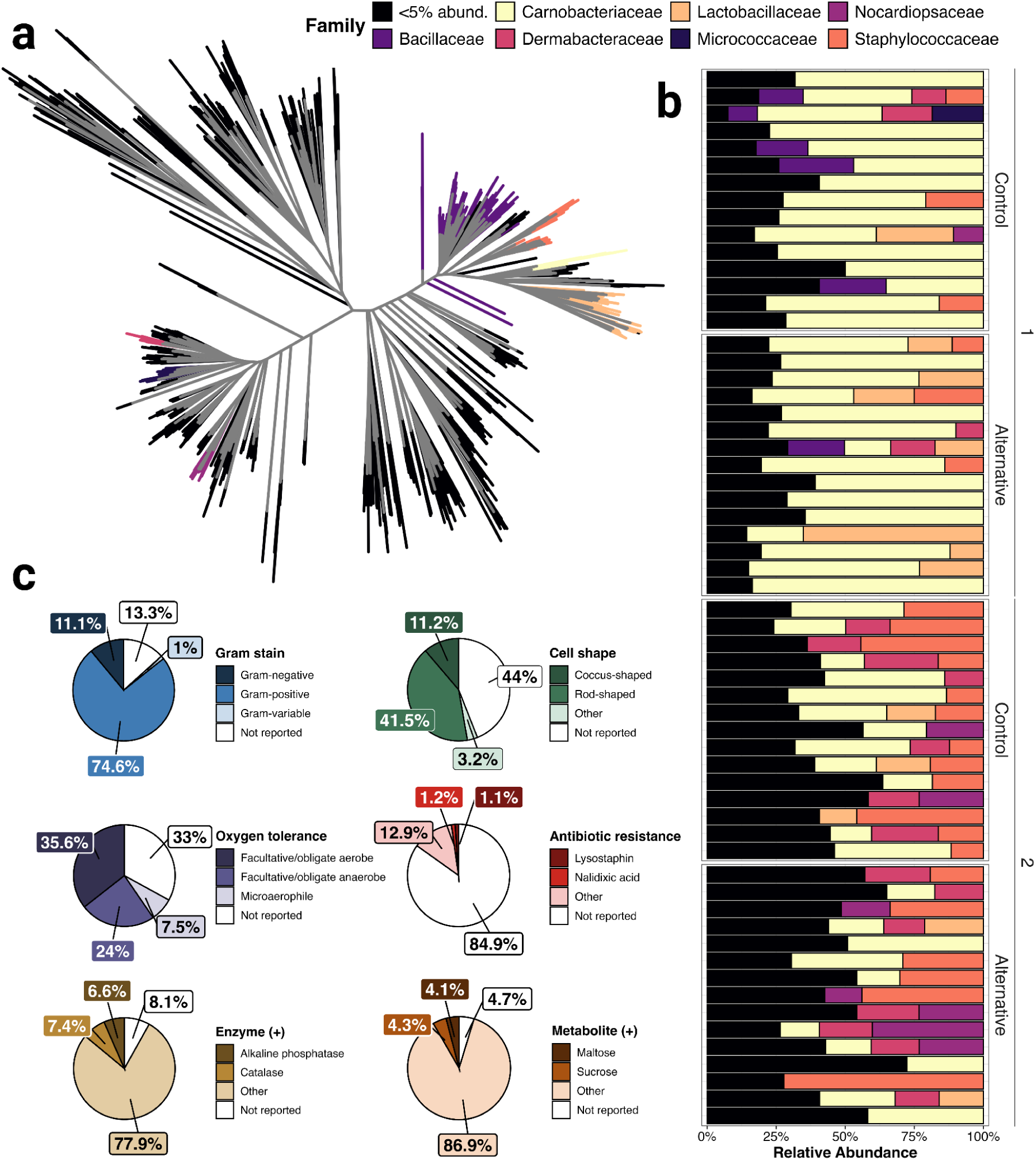
Microbiome overview. **(a)** Phylogenetic tree generated from MUSCLE-aligned 16S rRNA gene sequences. Maximum-likelihood sequence distances were calculated with phangorn, and the final tree was constructed using the neighbor-joining method, with branch support assessed across 1,000 bootstrap replicates. Tips represent features colored by family. **(b)** Stacked bar plot of family-level sample composition. Row facets reflect treatment and growout. **(c)** Pie charts of BacDive trait annotations.

Furthermore, cell morphologies were predominantly rods (41.5%) and cocci (11.2%), with the remainder unreported or less common forms. Oxygen tolerance was available for approximately two-thirds of species, with 35.6% aerobic, 24.0% anaerobic, and 7.5% microaerophilic. Known antibiotic resistance was noted for strains of <15% of detected species; among those, nalidixic acid (1.2%) and lysostaphin (1.1%) resistance were most frequently represented. The most utilized enzymes were catalase (7.4%) and alkaline phosphatase (6.6%), with the most utilized metabolites/substrates being sucrose (4.3%) and maltose (4.1%) (**Figure 2c**).

### α- and β-diversity varied with growout, house position, and alternative management

Treatment, growout, and position effects on α-diversity were determined using Shannon index, observed richness, inverse Simpson index, and Faith’s phylogenetic diversity. Growout significantly affected all metrics (Shannon, *p* = 1.06e-30; Observed richness, *p* = 2.38e-25; inverse Simpson, *p* = 5.38e-30; Faith’s PD, *p* = 7.12e-14). Position also showed global effects for Shannon (*p* = 0.004), inverse Simpson (*p* = 0.001), and Observed richness (*p* = 0.04). Within growout 1, position affected Shannon (*p* = 0.048), while within growout 2 it affected Shannon (*p* = 0.004) and inverse Simpson (*p* = 0.018). The treatment effect and the treatment-growout interaction were not detected at the global level (*p* > 0.05). However, pairwise contrasts within growout 2 indicated higher diversity in alternatively managed litter vs. Control for Faith’s PD (*p* = 0.04) and Observed richness (*p* = 0.02), with no treatment differences within growout 1 for any metric (**Figure 3a**). Adjusted intraclass correlation coefficients (ICCs) indicated <0.001% between-house variability for all α-diversity metrics in growout 1. In growout 2, between-house variability was 24.8% for observed richness, 6.7% for Shannon, <0.001% for inverse Simpson, and 33.6% for Faith’s PD. In the global models, adjusted ICCs were <0.01% for observed richness, Shannon, and inverse Simpson and 10.8% for Faith’s PD (**Supplementary File 2**).

**Figure 3.**
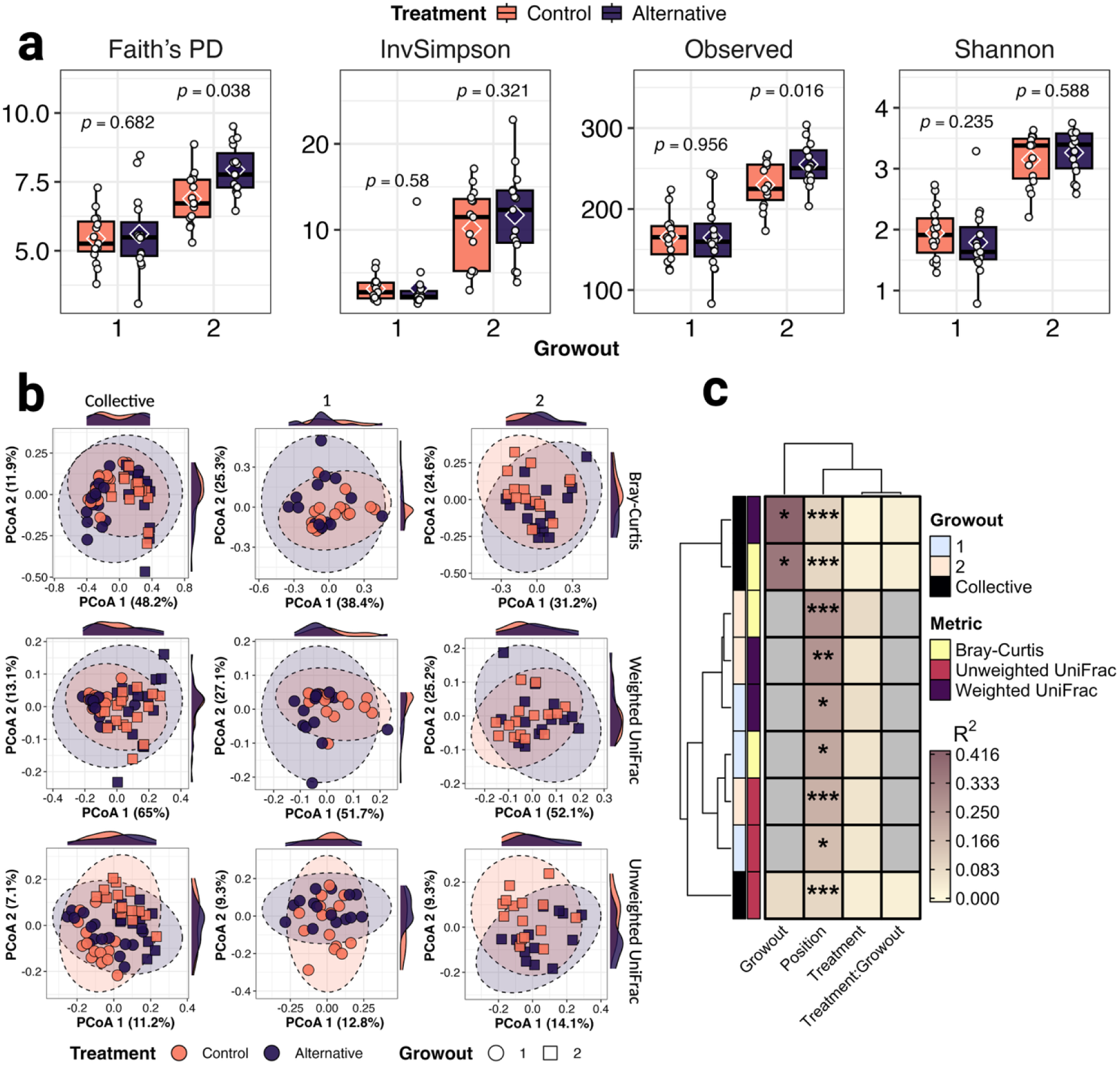
α and β diversity indices. **(a)** Mean estimates for Faith’s phylogenetic diversity (PD), inverse Simpson, observed richness, and Shannon diversity inferred across 1,000 subsampling iterations. **(b)** Principal Coordinates Analysis (PCoA) ordinations for Bray-Curtis, Weighted, and Unweighted UniFrac distances spanning collective and growout-specific analyses. **(c)** Heatmap depicting PERMANOVA partial R² values. Row annotations represent growout and distance metric. \*\*\**p* ≤ 0.001, \*\**p* ≤ 0.01, \**p* ≤ 0.05.

Treatment, growout, and position effects on β-diversity were assessed across Bray-Curtis, unweighted UniFrac, and weighted UniFrac dissimilarities by PERMANOVA with restricted permutations reflecting the experimental design. In the collective dataset, growout significantly explained variation for Bray-Curtis (*p* = 0.046, R² = 0.36) and weighted UniFrac (*p* = 0.046, R² = 0.42), but not unweighted UniFrac (*p* = 0.231, R² = 0.08). Position explained significant variation across all three metrics (Bray-Curtis, *p* = 3.0e-04, R² = 0.09; unweighted UniFrac, *p* = 4.0e-04, R² = 0.09; weighted UniFrac, *p* = 8.0e-04, R² = 0.10). Position remained significant within both growouts, explaining 15.6– 24.5% of variation in growout 1 (*p* = 0.011–0.037) and 19.2–28.5% in growout 2 (*p* = 2.0e-04–0.0097). Because treatment was assigned at the house level with three houses per treatment, the whole-house permutation space provided a minimum attainable treatment *p*-value of 0.10. Collective treatment effects explained 1.2–2.5% of variation across the three metrics (*p* = 0.10–0.20), while treatment-growout interactions explained approximately 2.0–2.3% (*p* = 0.10–0.30). Within growout 1, treatment explained 4.5–6.9% of variation (*p* = 0.20–0.30), compared with 5.0–8.0% within growout 2 (*p* = 0.10–0.20) (**Figure 3b**-**c**).

### Alternative management was associated with lower *Enterococcus* relative abundance and variable *in vitro* postbiotic responses

We next leveraged CPLM, NEGBIN, and ZINB models to identify differentially abundant taxa (*q* ≤ 0.05) between treatments, detecting 68 species across all models. Of these, 18 were identified in growout 1 comparisons (six enriched, 12 depleted in alternatively managed litter), 48 in growout 2 (18 enriched, 30 depleted), and 27 in the collective dataset (eight enriched, 19 depleted). All species shared across time-stratified models showed consistent directional changes, including five shared between growouts 1 and 2, seven shared between growout 1 and the collective dataset models, and 18 shared between growout 2 and the collective dataset models. Among differentially abundant species, *Brevibacterium anseongense* and *B. profundi* were each detected in the most models (*n* = 14) and were consistently depleted under alternative management. The next most frequently detected species, *B. avium* (*n* = 12), was likewise depleted (**Figure 4a**, **Supplementary Figure 3**).

**Figure 4.**
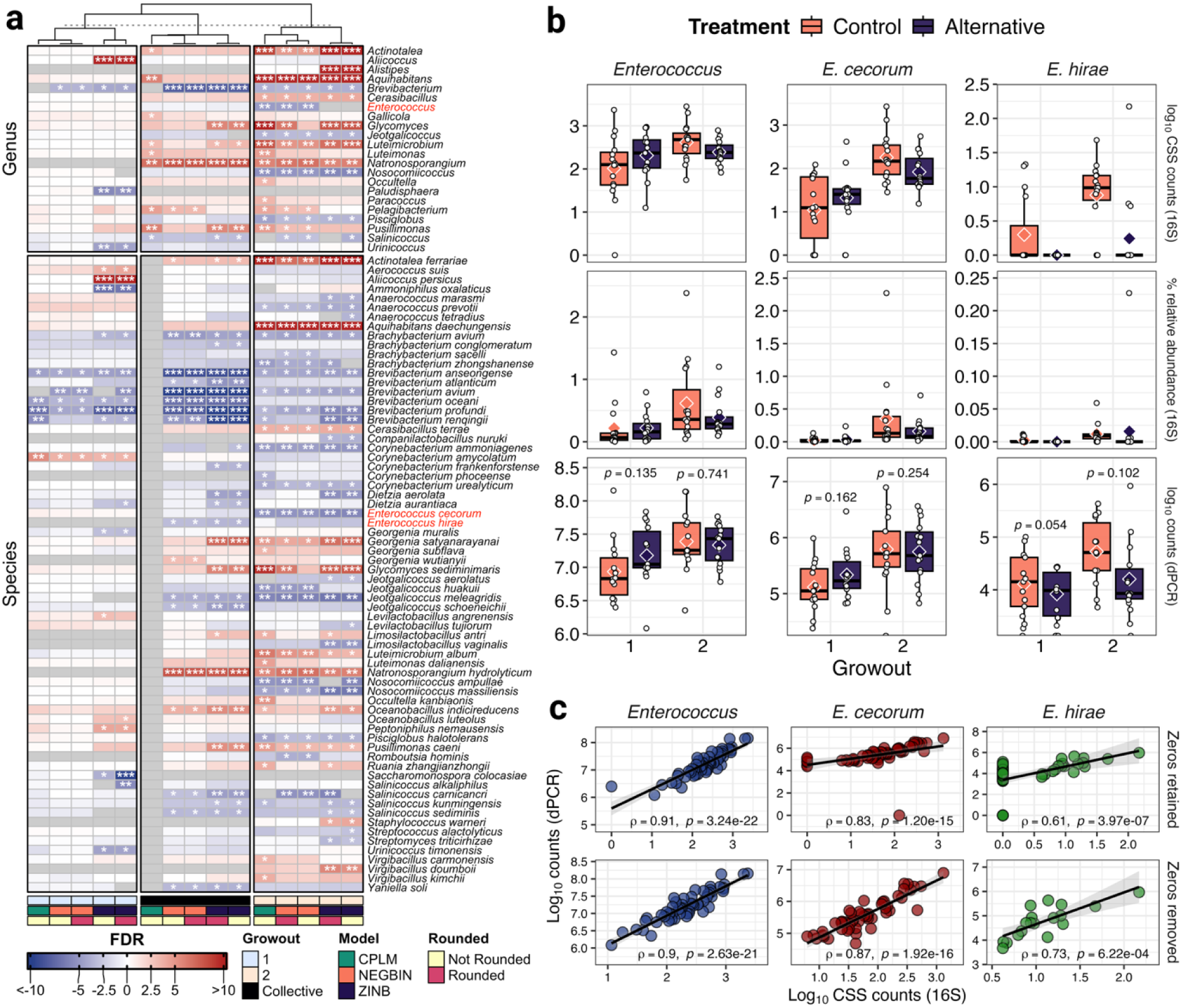
Differential abundance analysis. **(a)** Heatmap depicting differentially abundant features between treatments (*q* ≤ 0.05). Column annotations reflect growout, statistical model, and data rounding, while row partition reflects analyses conducted at species and genus levels. Cell color represents enrichment (red) or depletion (blue) in alternatively managed litter vs. Control. \*\*\**q* ≤ 0.001, \*\**q* ≤ 0.01, \**q* ≤ 0.05. (**b**) Sequencing- and dPCR-based abundance patterns for *Enterococcus*, *E. cecorum*, and *E. hirae*. Boxplots show log₁₀ CSS-normalized 16S counts, percent relative abundance from 16S data, and log₁₀ dPCR copy number estimates. (**c**) Correlations between sequencing-derived log₁₀ CSS-normalized counts and log₁₀ dPCR counts for *Enterococcus*, *E. cecorum*, and *E. hirae*. Associations are shown with zeros retained and after zero removal.

At genus level, we identified 22 genera that were differentially abundant in at least one model. Four genera were detected in growout 1 (one enriched, three depleted), 18 in growout 2 (12 enriched, six depleted), and 11 in the collective dataset (nine enriched, two depleted). Consistent with species-level patterns, all genera shared across datasets showed the same directional changes, including one shared between growouts 1 and 2, one shared between growout 1 and the collective dataset, and ten shared between growout 2 and the collective dataset. *Brevibacterium* was detected in the most models (*n* = 13) and was consistently depleted, followed by *Natronosporangium* (*n* = 10; enriched) and *Luteimicrobium* (*n* = 8; enriched) (**Figure 4a**, **Supplementary Figure 3**).

Verification of species-level assignments showed that *E. cecorum* reads were well resolved from other *Enterococcus* spp. Although most *E. hirae* reads (80%) aligned most closely to *E. hirae*, none exceeded the closest alternative species by the 0.5 percentage-point resolution criterion, and sequencing-derived *E. hirae* abundance is therefore reported as the *E. hirae* group. Alternatively managed litter had significantly lower relative abundance of *Enterococcus cecorum* (growout 2 FDR < -5), *E. hirae* group (collective FDR < -3), and genus-level *Enterococcus* (growout 2 FDR < -4.5) (**Figure 4a**). Genus- and species-level dPCR assays were therefore developed to determine whether these relative abundance shifts reflected changes in absolute abundance. The relative and absolute abundances were strongly correlated in all instances (*Enterococcus*, ρ = 0.91, *p* = 3.24e-22; *E. cecorum*, ρ = 0.83, *p* = 1.2e-15; *E. hirae*, ρ = 0.61, *p* = 3.97e-07), with improved correspondence at species level following exclusion of samples lacking 16S detection (*E. cecorum*, ρ = 0.87, *p* = 1.92e-16; *E. hirae*, ρ = 0.73, *p* = 6.22e-04) (**Figure 4c**). At genus level, growout significantly influenced *Enterococcus* abundance (*p* = 0.006), whereas treatment (*p* = 0.40), position (*p* = 0.53), and the treatment-growout interaction (*p* = 0.19) were not detected. Consistent with these results, no treatment differences were observed within either growout based on pairwise contrasts (*p* ≥ 0.14), while position significantly affected *Enterococcus* abundance within growout 2 (*p* = 5.48e-06) but not growout 1 (*p* = 0.12) (**Figure 4b**). At species level, *E. cecorum* abundance was significantly influenced by growout (*p* = 9.11e-07) and position (*p* = 1.15e-04), with a marginal treatment-growout interaction (*p* = 0.07), whereas treatment was not detected globally (*p* = 0.77) or within growouts (*p* ≥ 0.16). In contrast, *E. hirae* exhibited a significant global treatment effect (*p* = 0.01), corresponding to lower absolute abundance overall in alternatively managed litter. This pattern is consistent with its lower relative abundance in the *MaAsLin2* models for the collective dataset. Growout (*p* = 3.95e-05) and position (*p* = 8.42e-04) also significantly influenced *E. hirae* abundance. However, treatment differences were not detected within growouts (*p* ≥ 0.05). Position effects were again strongest within growout 2 for both species (*E. cecorum*, *p* = 0.002; *E. hirae*, *p* = 9.17e-05), with significant but weaker effects observed in growout 1 (*E. cecorum*, *p* = 0.004; *E. hirae*, *p* = 0.04) (**Figure 4b**). House-level variability was negligible in the global GLMMs, with adjusted ICCs <0.001% for genus-level *Enterococcus*, *E. cecorum*, and *E. hirae*. Greater between-house variability was observed in growout 1 for genus-level *Enterococcus* (16.6%) and *E. hirae* (32.5%), while all other growout-specific ICCs were <0.001% (**Supplementary File 2**).

To characterize *in vitro* responses of *Enterococcus* to the postbiotic fraction of the alternative management strategy, we performed 24-h adherent-biomass assays using *E. cecorum* and *E. hirae* ATCC isolates across a concentration gradient of the postbiotic solution (**Figure 5a**). Assays included an untreated bacterial control (BCT) and vancomycin (8 µg/mL) as a positive control. Treatment significantly affected adherent biomass for both *E. cecorum* (ANOVA *p* = 1.92e-13) and *E. hirae* (*p* = 2.48e-05). For *E. cecorum*, adherent biomass was 8.80 to 13.80% lower than BCT at concentrations ≤6%, increasing to 25.10% at 12% and 36.80% at 24% (**Figure 5b**). Pairwise comparisons with BCT were significant at 1.5, 6, 12, and 24% (*p* ≤ 0.02), but not at 0.75 or 3%, while the 12% and 24% treatments also differed from lower concentrations (*p* ≤ 0.03). *E. hirae* exhibited a comparatively flatter response, with adherent biomass 9.00 to 15.60% lower than BCT across postbiotic concentrations (**Figure 5b**). Differences from BCT were detected at 0.75, 3, 12, and 24% (*p* ≤ 1.61e-03), but not at 1.5 or 6%, while differences among postbiotic concentrations were generally not detected. At 6%, within the nominal application-mixture range (5.7–9.1%), adherent biomass was 13.80% lower than BCT for *E. cecorum* and 9.40% lower for *E. hirae*, with only the *E. cecorum* response differing significantly from BCT. Overall, responses near the nominal application-mixture range were modest and not consistently detected across both species, whereas the largest reductions in adherent biomass occurred at higher postbiotic concentrations.

**Figure 5.**
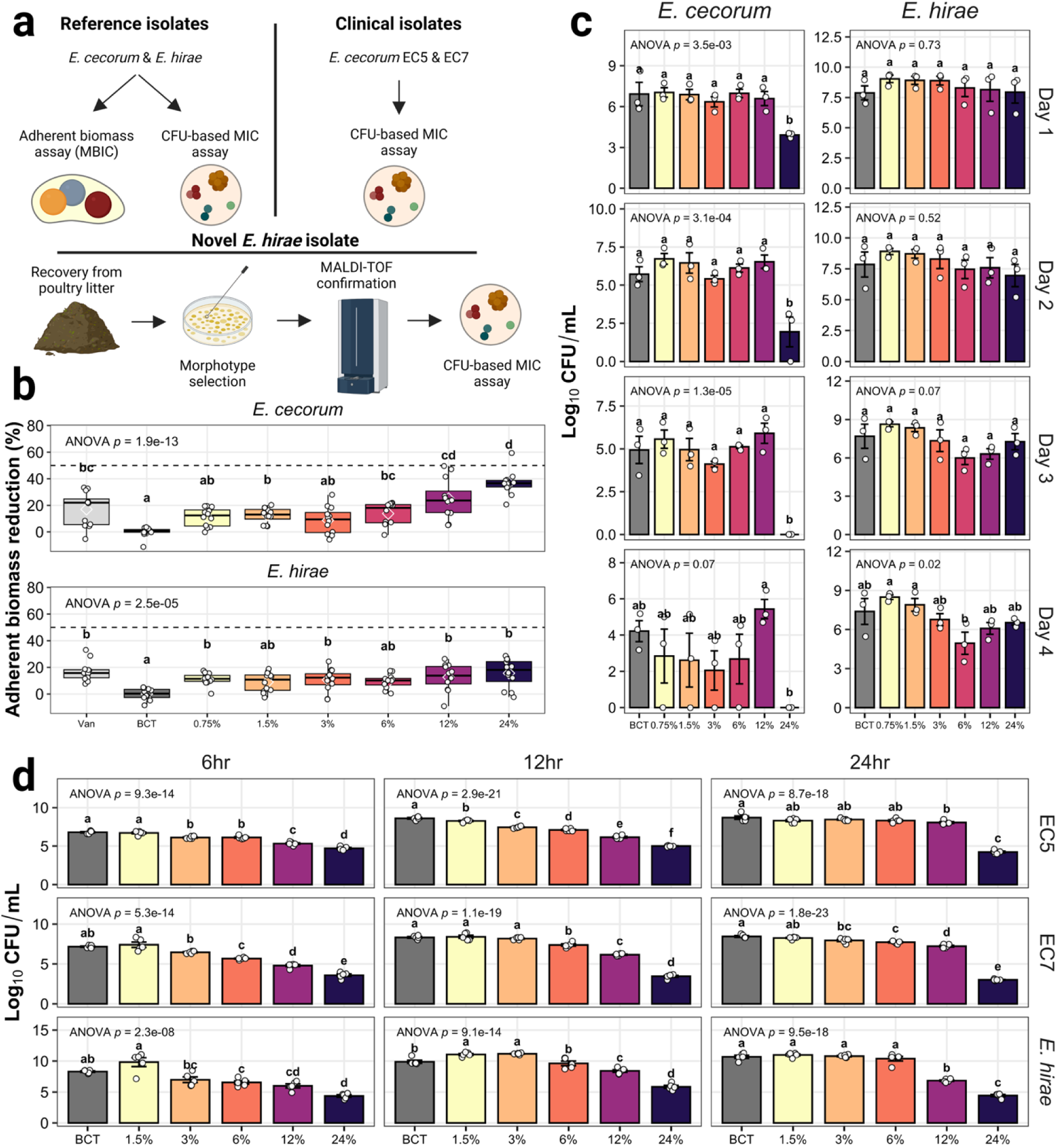
In vitro responses of *Enterococcus* isolates to the postbiotic fraction. **(a)** Schematic overview of adherent-biomass and CFU-based MIC assays performed with ATCC reference isolates, clinical *E. cecorum* isolates, and a litter-derived *E. hirae* isolate. **(b)** Adherent-biomass assay results for ATCC *E. cecorum* and *E. hirae* using a conventional minimum biofilm inhibitory concentration (MBIC) framework. The dashed line indicates a 50% reduction in adherent biomass relative to the untreated bacterial control (BCT). **(c)** CFU-based MIC assays for ATCC *E. cecorum* and *E. hirae* across four days of exposure. **(d)** CFU-based MIC assays for clinical *E. cecorum* isolates EC5 and EC7 and the litter-derived *E. hirae* isolate across 6, 12, and 24 h of exposure.

We next evaluated postbiotic-associated changes in viable recovery using CFU-based MIC assays across four timepoints (**Figure 5c**). Treatment significantly influenced *E. cecorum* recovery on days 1 to 3 (ANOVA *p* ≤ 3.46e-03), with weaker evidence by day 4 (*p* = 0.07) (**Figure 5c**). The clearest response occurred at 24%, corresponding to 3.01-, 3.78-, and 4.94-log₁₀ lower recovery relative to BCT on days 1–3, respectively, with no detectable colonies from day 3 onward. These differences were significant in pairwise comparisons on days 1–3 (*p* ≤ 6.30e-03) (**Figure 5c**). At 6%, *E. cecorum* recovery remained statistically indistinguishable from BCT across all timepoints, with mean recovery ranging from 0.05-log₁₀ lower to 0.40-log₁₀ higher than BCT through day 3 and 1.54-log₁₀ lower by day 4 (**Figure 5c**). For *E. hirae*, an overall treatment effect was detected only on day 4 (*p* = 0.02) (**Figure 5c**). The largest numerical difference from BCT occurred at 6%, reaching 2.44-log₁₀ lower recovery by day 4; however, no individual postbiotic concentration differed significantly from BCT in pairwise comparisons at any timepoint (*p* ≥ 0.12) (**Figure 5c**). At lower concentrations (≤1.5%), CFU recovery was frequently similar to or greater than BCT, particularly for *E. hirae*. Overall, the CFU-based MIC assays showed substantial concentration-and species-dependent variability, with statistically supported reductions in viable recovery resolved only for *E. cecorum* at 24%.

To determine whether the ATCC strain patterns extended to broiler-derived clinical isolates, we evaluated EC7 and EC5, two published *E. cecorum* isolates recovered from diseased broilers, using a truncated postbiotic concentration gradient from 1.5 to 24% over 6, 12, and 24 h (**Figure 5a**, **d**). Treatment significantly affected recovery for both isolates at all timepoints (ANOVA EC7, *p* ≤ 1.21e-23; EC5, *p* ≤ 2.33e-29). For EC7, recovery at 6% was 1.49-, 0.95-, and 0.74-log₁₀ lower than BCT at 6, 12, and 24 h, respectively (*p* ≤ 1.96e-04) (**Figure 5d**), with the magnitude of the difference decreasing over time. At 12%, EC7 recovery was 1.22- to 2.38-log₁₀ lower than BCT, while at 24% it was 3.59- to 5.45-log₁₀ lower (*p* ≤ 2.06e-08) (**Figure 5d**). EC5 showed a less persistent response at 6%, with 0.68- and 1.51-log₁₀ lower recovery at 6 and 12 h, respectively (*p* ≤ 4.45e-04), but no difference from BCT by 24 h (*p* = 0.33) (**Figure 5d**). At 12%, EC5 recovery was 0.65- to 2.47-log₁₀ lower than BCT (*p* ≤ 2.21e-02), while 24% produced 2.09- to 4.47-log₁₀ lower recovery across timepoints (*p* ≤ 6.50e-13) (**Figure 5d**). Thus, responses at 6% were isolate- and time-dependent and diminished with continued exposure, whereas larger and more persistent differences in viable recovery were observed at 12 and 24%.

We next recovered an *E. hirae* isolate from poultry litter, identified it by MALDI-TOF (data not shown), and tested it as a study-derived comparator to the ATCC *E. hirae* isolate (**Figure 5a**, **d**). Treatment influenced recovery at 6, 12, and 24 h (ANOVA *p* ≤ 1.31e-11), but reductions were largely restricted to higher postbiotic concentrations. At 6%, recovery was 1.76-log₁₀ lower than BCT at 6 h (*p* = 0.0465), but this effect was not maintained at 12 or 24 h (*p* ≥ 0.91) (**Figure 5d**). More consistent differences were observed at 12% and 24%, with 12% postbiotic producing 1.49 to 3.85-log₁₀ lower recovery and 24% producing 3.94 to 6.26-log₁₀ lower recovery across timepoints (*p* ≤ 5.00e-03) (**Figure 5d**). In contrast, 1.5% and 3% postbiotic increased *E. hirae* recovery at 12 h relative to BCT by 1.19- and 1.29-log₁₀, respectively (*p* ≤ 0.02).

### Environmental gradients were associated with prokaryotic microbiome structure

The litter physicochemical profile (NH₄⁺, total N, P, K, and moisture) was quantified and compared between treatments, growouts, and positions on a dry-weight basis. Composite baseline samples showed no significant differences between treatments for NH₄⁺, P, or K prior to study initiation (*p* > 0.05), with total N and moisture being marginally elevated in Alternative houses (*p* = 0.05). These trends largely persisted in growout 1, with total N (*p* = 0.01), P (*p* = 0.02), and moisture (*p* = 0.03) being higher in Alternative. Interestingly, P and K showed an inverse pattern in growout 2, being significantly elevated in the Control (*p* = 0.04 and 3.86e-04, respectively). Moisture and total N stabilized across treatments, with the percent change relative to the baseline mean being 5.24% and 27.01% for Alternative, and 39.45% and 78.16% for the Control, respectively. In the global GLMMs, treatment had no significant main effects for any metric, whereas growout significantly affected total N (*p* = 0.01), NH₄⁺/total N (*p* = 0.01), P (*p* = 3.48e-13), K (*p* = 2.81e-06), and moisture (*p* = 0.006). Position significantly affected NH₄⁺ (*p* = 0.037), total N (*p* = 0.012), P (*p* = 0.001), K (*p* = 0.006), and moisture (*p* = 1.69e-06). Within growout 1, position affected total N, P, K, and moisture (*p* ≤ 0.002), whereas within growout 2 it affected NH₄⁺, NH₄⁺/total N, and moisture (*p* ≤ 0.043). Significant treatment-growout interactions were detected for total N (*p* = 0.02), P (*p* = 2.14e-03), K (*p* = 1.62e-04), and moisture (*p* = 1.50e-03) (**Figure 6a**). Adjusted ICCs were <0.001% in the global models for NH₄⁺, total N, NH₄⁺/total N, P, and K, compared with 16.6% for moisture. Within growout 1, the largest house-level ICCs were observed for NH₄⁺/total N (42.8%), moisture (21.0%), and NH₄⁺ (16.9%), with smaller values for P (9.8%) and K (3.8%) and <0.001% for total N. In growout 2, the ICC was 25.0% for moisture and 1.2% for total N, while all remaining metrics were <0.001% (**Supplementary File 2**).

**Figure 6.**
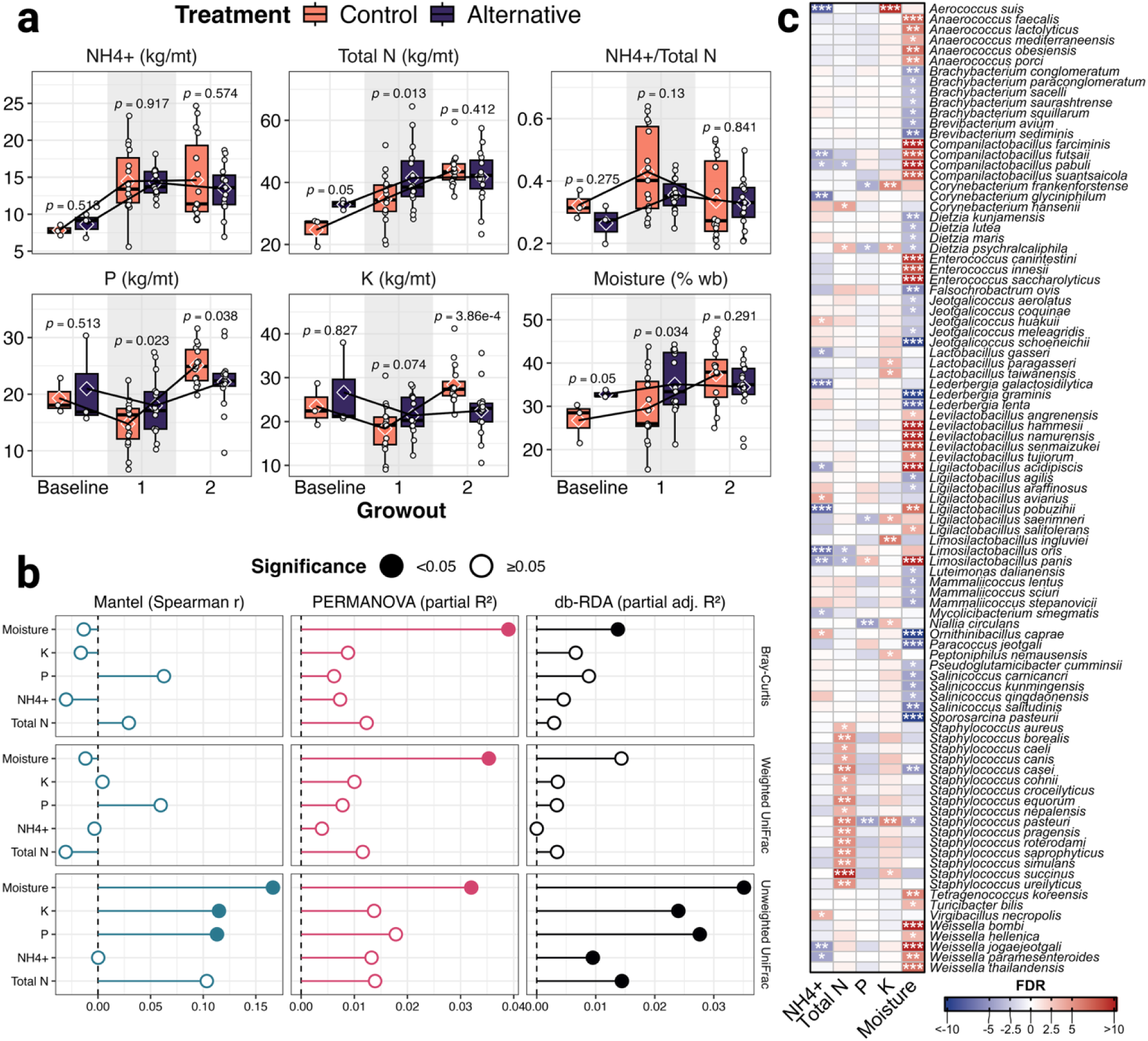
Physicochemical analysis and correlation with microbiome parameters.(a) Physicochemical metrics spanning a baseline (prior to study initiation) and growouts 1 and 2. **(b)** Effect of physicochemical metrics on microbiome composition. Inferences were drawn with Mantel, PERMANOVA, and db-RDA analyses, and spanning Bray-Curtis, Weighted, and Unweighted UniFrac distances. **(c)** Heatmap depicting features significantly associated with one or more physicochemical metrics (*q* < 0.05). \*\*\**q* ≤ 0.001, \*\**q* ≤ 0.01, \**q* ≤ 0.05.

Relationships between microbial community structure and the physicochemical environment were evaluated using Bray-Curtis, unweighted UniFrac, and weighted UniFrac dissimilarity matrices. Overall correspondence between each distance metric and the standardized environmental matrix (N, NH₄⁺, P, K, and moisture) was assessed using Procrustean randomization tests. Variable-specific associations were then evaluated using Mantel tests. Neither Bray-Curtis nor weighted UniFrac dissimilarity was significantly correlated with any individual physicochemical variable. Unweighted UniFrac dissimilarity was significantly correlated with moisture (*r* = 0.17, *p* = 0.004), with a marginal association observed for K (*r* = 0.12, *p* = 0.056) (**Figure 6b**).

To assess associations between environmental parameters and β diversity after adjusting for experimental structure, physicochemical variables were incorporated as covariates in PERMANOVA models. Among the environmental variables examined, moisture was associated with Bray-Curtis (R² = 0.03, *p* = 0.003) and unweighted UniFrac (R² = 0.03, *p* = 8.0e-04), with a weaker, marginally significant association for weighted UniFrac (R² = 0.02, *p* = 0.064). P was additionally associated with unweighted UniFrac (R² = 0.02, *p* = 0.045). Total N, NH₄⁺, and K were not significantly associated with β diversity for any distance metric. These results indicate that, after accounting for primary experimental factors, litter moisture was the most consistent physicochemical correlate of between-sample microbial community variation (**Figure 6b**).

To further evaluate the contribution of individual physicochemical variables to β diversity, we applied partial db-RDA models that accounted for house, growout, treatment, and position prior to testing each environmental constraint. For Bray-Curtis, moisture was the only significant physicochemical variable (partial adj. R² = 0.014, *p* = 0.026). No physicochemical variable was significantly associated with weighted UniFrac. In contrast, unweighted UniFrac was significantly associated with total N (partial adj. R² = 0.014, *p* = 0.007), NH₄⁺ (partial adj. R² = 0.010, *p* = 9.0e-04), P (partial adj. R² = 0.028, *p* = 1.0e-04), K (partial adj. R² = 0.024, *p* = 1.0e-04), and moisture (partial adj. R² = 0.035, *p* = 1.0e-04) (**Figure 6b**).

Lastly, we employed a compound Poisson linear model to identify taxa significantly associated (*q* < 0.05) with each environmental parameter, detecting a collective 117 taxa. Across parameters, moisture yielded the greatest number of associations (*n* = 62 taxa; 28 positive, 34 negative), followed by total N (*n* = 21; 18 positive, 3 negative), NH₄⁺ (*n* = 17; 4 positive, 13 negative), K (*n* = 11; all positive), and P (*n* = 6; 1 positive, 5 negative). Several taxa exhibited multi-parameter associations. *Limosilactobacillus panis* was positively associated with moisture and P but negatively associated with NH₄⁺ and total N. *Staphylococcus pasteuri* showed positive associations with total N and K alongside negative associations with moisture and P, and *Dietzia psychralcaliphila* displayed a similar four-parameter profile, with positive associations with total N and K and negative associations with P and moisture. At the genus level, the strongest single-parameter pattern was observed for *Staphylococcus*, with 16 taxa positively associated with total N. Additional genus-level patterns were evident for moisture, including *Anaerococcus* (five species; all positive), *Levilactobacillus* (five species; all positive), *Weissella* (five species; all positive), and *Companilactobacillus* (four species; all positive), whereas *Brachybacterium* (five species), *Dietzia* (four species), *Jeotgalicoccus* (four species), and *Salinicoccus* (four species) were all negatively associated with moisture. *Enterococcus* comprised three species that were all positively associated with moisture (**Figure 6c**).

## Discussion

### Full-length 16S rRNA gene sequencing captured the anticipated prokaryotic community

This study evaluated how an alternative litter management regimen (postbiotic intervention plus half-rate NaHSO₄) influences prokaryotic community assembly, *Enterococcus* abundance, and litter physicochemistry in a commercial broiler rearing system. Microbiome composition inferred from full-length 16S sequencing was taxonomically consistent with prior reports, with Firmicutes as the dominant phylum followed by Actinobacteria [76–78]. This taxonomic structure corresponded with expected physiological traits, as BacDive annotation indicated a community composed predominantly of Gram-positive organisms (∼75%). Poultry litter microbiomes are Gram-positive dominated, with estimates ranging from 50 to 88% based on culture- and sequencing-based approaches [12, 77, 79–81]. Furthermore, taxa identified in this study were primarily rod- or coccus-shaped and encompassed a heterogeneous mix of aerobic and anaerobic organisms relative to proportions reported for the broiler gastrointestinal tract [82]. These patterns collectively indicate that the dataset captured the expected structure of the broiler litter microbiome, providing a reliable foundation for evaluating shifts in community assembly under alternative litter management.

### Alternative litter management influenced microbiome structure across space and time

Broad treatment effects on microbiome structure and composition were evaluated using α- and β-diversity metrics. Treatment-associated differences were most evident in growout 2, after two consecutive flocks under alternative management. These included higher observed richness and phylogenetic diversity, while β-diversity differences remained modest but accounted for more variation than in growout 1. This pattern is consistent with Olson et al. [21], which reported greater compositional divergence and more differentially abundant taxa following consecutive alternative-management exposures relative to a single exposure.

Spatial position and growout were major sources of variation in litter microbiome structure and physicochemistry and were generally associated with greater effects than management, indicating that treatment responses occurred within a heterogeneous and maturing litter environment. Positional effects were observed for three of four α-diversity metrics and five physicochemical parameters, necessitating inclusion of position as a covariate in β-diversity and differential abundance models. This is consistent with prior work linking within-house heterogeneity to microclimate variation and litter microbiome structure [5, 24, 83]. The spatially aware sampling framework used here is conceptually similar to zone-based soil sampling, where known spatial variability is represented by subdividing the environment into discrete units [84]. Temporal effects were similarly evident across diversity, composition, and physicochemistry, with growout 2 exhibiting increased diversity and greater compositional dissimilarity compared to growout 1. Together, these patterns indicate progressive community maturation across successive flocks, consistent with longitudinal observations that temporal dynamics can outweigh individual management effects once litter communities are established [19]. They also emphasize the importance of reporting litter age, management history, and spatial sampling design in poultry microbiome studies.

### Management-associated shifts in *Enterococcus* are consistent with broader litter community restructuring

Management practices applied during downtimes are critical in shaping poultry litter microbiomes, where competitive exclusion influences pathogen persistence and the emergence of disease-suppressive environments [18]. Previous commercial and microcosm studies showed selective and context-dependent microbial responses to postbiotic-based regimens, including treatment-associated differences among presumptive enterococci and staphylococci and lower *Staphylococcus* relative abundance under commercial conditions [20–21]. Such patterns were observed in the present study, where alternatively managed litter exhibited significantly lower relative abundance of total *Enterococcus* and *E. cecorum* in growout 2, alongside lower relative abundance of *E. hirae* group across the collective dataset. These taxa include well-established facultative pathogens in poultry [85], making the observed compositional shifts potentially relevant to flock health. Absolute quantification by dPCR was correlated with sequencing-based abundance patterns across samples, although treatment-associated lower relative abundance detected by 16S was not consistently resolved as lower absolute copy number. This suggests that alternative management altered the proportional dominance of *Enterococcus* within the litter microbiome without producing uniform reductions in absolute abundance, potentially through broader changes in the litter environment and microbial community structure. A nonexclusive explanation is that management-associated changes altered community assembly and relative competitiveness, while absolute abundance remained sufficiently variable by terminal sampling to obscure corresponding treatment effects by dPCR. Additionally, because both 16S rRNA gene sequencing and standard dPCR quantify DNA rather than viable cells, relic DNA from nonviable *Enterococcus* cells may have contributed to endpoint copy number estimates. Under these scenarios, relative abundance profiles would retain evidence of compositional displacement even when endpoint absolute quantification does not show a corresponding uniform reduction in abundance.

These overlapping suppositions are further informed by *in vitro* response assays, which showed species-, strain-, concentration-, and exposure-time-dependent responses to the postbiotic. At 6%, the postbiotic produced modest reductions in adherent biomass, whereas CFU-based responses at the same concentration were weaker, transient, or not consistently different from BCT. This divergence between adherent biomass and viable recovery suggests that responses near the nominal application-mixture range were limited and phenotype-dependent rather than indicative of uniform changes in viability. Greater reductions were observed at concentrations above 6%, particularly for ATCC *E. cecorum* and the EC5 and EC7 clinical isolates. In contrast, ATCC and study-derived *E. hirae* isolates showed less consistent responses at lower exposure levels, including increased recovery of the poultry litter isolate at 1.5 to 3% postbiotic. These strain-variable responses align with documented heterogeneity among poultry-associated *Enterococcus* spp. [85] and with prior evidence that microbial metabolites can affect pathogenic *E. cecorum* in an isolate-dependent manner, with responses ranging from complete suppression to growth promotion [86]. Collectively, the variable magnitude and persistence of these responses suggest that antimicrobial activity alone is unlikely to account for the *Enterococcus* relative abundance patterns observed under commercial conditions.

Observations herein are consistent with a model in which shifts in *E. cecorum* and *E. hirae* group relative abundance occur within broader restructuring of the litter microbiome, with lower proportional dominance of these taxa occurring alongside changes in competitive, commensal, or mutualistic populations. In the broiler gastrointestinal tract, cecal fermentation broth reduced *Enterococcus* and *Escherichia-Shigella* while increasing *Bacteroides* and Ruminococcaceae [87], illustrating that suppression of facultative opportunists can coincide with expansion of taxa associated with competitive exclusion. These community-level shifts may also be reinforced by the physicochemical component of alternative management, since half-rate NaHSO₄ may transiently alter pH and microbial dynamics while reducing salt input, potentially disadvantaging salt-tolerant enterococci relative to other litter-adapted taxa [88]. Future work should disentangle these mechanisms by combining controlled microcosms, strain-resolved metagenomics, and metabolomic or ionomic profiling to resolve resource competition, niche occupancy, and physicochemical drivers of community restructuring.

### Integrated microbiome monitoring can guide poultry litter management

This study provides a foundation for connecting poultry litter microbiome structure to measurable environmental and management variables under commercial conditions. By integrating full-length 16S sequencing with physicochemistry, we observed consistent shifts in community composition and environmental parameters across successive growouts, indicating that litter maturation is accompanied by reproducible changes in microbial assembly. The incorporation of spatial variability into both sampling and statistical analyses further demonstrates that within-house heterogeneity can be captured and should be considered when interpreting litter microbiome data or developing predictive models [89]. The addition of nanoplate dPCR extends this framework by enabling absolute quantification of selected taxa in poultry litter, an inhibitor-rich matrix where dPCR offers advantages over standard curve-dependent qPCR [28–29, 90]. Paired with sequencing, this approach distinguished proportional shifts in *Enterococcus* from changes in absolute abundance, strengthening interpretation of management-associated community restructuring. Adherent biomass and CFU-based MIC assays provided complementary context for these patterns, revealing concentration-, strain-, and exposure-dependent responses. The shifts in diversity and taxon dominance observed under alternative management illustrate how integrated microbiome monitoring can evaluate management-associated responses and inform litter management, reuse, and nutrient management across production cycles.

### Study limitations

Limitations of the present study should be considered in the context of integrator policy and biosecurity constraints inherent to commercial broiler production. First, treatment-level replication was limited to three independent houses per management regimen. Because house was the true experimental unit, the resulting whole-house permutation space for β-diversity yielded a minimum attainable treatment *p*-value of 0.10, limiting formal inference regarding treatment effects on community composition. Second, a true baseline microbiome was not established prior to treatment implementation, limiting the ability to resolve pre-existing differences between houses and to attribute observed shifts exclusively to management. Nevertheless, comparable physicochemical profiles and production history across houses suggest similar starting conditions between Control and Alternative regimens. Third, microbiome characterization was performed at a single terminal timepoint per growout, whereas broiler litter microbial communities are known to undergo dynamic shifts throughout the production cycle [78, 91]. As such, the present dataset captures end-point community structure but does not resolve within-growout temporal trajectories or transient responses to treatment. Lastly, the alternative management regimen combined postbiotic administration with reduced NaHSO₄ application, precluding direct attribution of observed microbiome and physicochemical shifts to individual components. While postbiotic-associated responses of *Enterococcus* are supported by prior microcosm findings [20], other patterns, including lower relative abundance of halotolerant taxa (e.g., *Brevibacterium* spp.), are more consistent with reduced NaHSO₄ inputs [92]. Evaluation of reduced NaHSO₄ application in commercial systems also requires direct assessment of NH₃ and, ideally, litter pH dynamics, which were not measured in this study. Although NH₃ was monitored by the integrator and no issues were reported during the trial, this may not be representative of all commercial broiler operations. Disentangling the relative contributions of biologic and chemical drivers and establishing the feasibility of reduced acidifying inputs across production systems will require factorial experimental designs that independently vary amendment components while incorporating continuous NH₃ and physicochemical monitoring.

## Conclusion

These findings demonstrate that alternative litter management with a postbiotic intervention and reduced-rate NaHSO₄ influenced poultry litter prokaryotic community structure under commercial broiler production conditions. Treatment-associated differences were most evident in observed richness and phylogenetic diversity, lower relative abundance of *Enterococcus* spp. (including *E. cecorum* and *E. hirae* group), and modest variation in community composition, while position, litter age, moisture, and nutrient gradients remained important sources of variation. Full-length 16S rRNA gene sequencing, dPCR, and isolate-level assays indicate that *Enterococcus* responses are more consistent with changes in proportional dominance and broader community restructuring than with uniform reductions in absolute abundance. Future and ongoing efforts should incorporate higher-resolution temporal profiling, NH₃ and pH monitoring, quantification of antimicrobial resistance and virulence gene content, and controlled trials that elucidate the relative contributions of postbiotic and acidifier components to prokaryotic litter microbiome restructuring. These approaches will help define how litter amendments influence microbial ecology and broiler health under commercial production conditions.

## Abbreviations

NH₃: ammonia; NaHSO₄: sodium bisulfate; dPCR: digital PCR; ONT: Oxford Nanopore Technologies; MSA: multiple sequence alignment; UPGMA: Unweighted Pair Group Method with arithmetic mean; AIC: Akaike Information Criterion; GLMM: generalized linear mixed model; CSS: cumulative sum scaling; PcoA: Principal Coordinates Analysis; PERMANOVA: permutational multivariate analysis of variance; MaAsLin2: Microbiome Multivariable Associations with Linear Models; NEGBIN: negative binomial; ZINB: zero-inflated negative binomial; CPLM: compound poisson linear model; FDR: false discovery rate; FAM: 6-carboxyfluorescein; BHQ1: Black Hole Quencher 1; HEX: hexachloro-fluorescein; dMIQE: Minimum Information for Publication of Quantitative Digital PCR Experiments; LOB: limit of blank; LOD: limit of detection; LOQ: limit of quantification; MBIC: minimum biofilm inhibitory concentration; PBS: phosphate-buffered saline; TSB: tryptic soy broth; ATCC: American Type Culture Collection; CFU: colony-forming unit; HSD: Tukey’s honestly significant difference; MIC: minimum inhibitory concentration; BHI: brain heart infusion; MALDI-TOF: matrix-assisted laser desorption/ionization time-of-flight; N: nitrogen; NH₄⁺: ammonium-N; P: phosphorus; K: potassium; RMMA: Recommended Methods of Manure Analysis; wb: wet basis; PROTEST: Procrustean randomization test; db-RDA: partial distance-based redundancy analysis; PD: phylogenetic diversity; ICC: intraclass correlation coefficient

## Ethics approval and consent to participate

Not applicable

## Consent for publication

Not applicable

## Availability of data and materials

The raw 16S sequencing data have been deposited in the NCBI Sequence Read Archive under the accession number PRJNA1466458. The R and R Markdown files used for data analysis are available in the GitHub repository, https://github.com/brett-hale/Alternative-managment-strategy.

## Author contributions

B.H. conceived and designed the study, secured funding, collected samples, performed nucleic acid isolation, data analysis and visualization, and drafted the manuscript. C.P. collected samples and performed nucleic acid extraction and purification. G.G. performed dPCR assays. C.A. performed nucleic acid extraction and purification. K.T. and R.C.S. performed MIC assays with ATCC isolates. L.D. and R.C.S. performed MBIC assays with ATCC isolates. M.C. and D.G. performed MIC assays with EC5 and EC7 isolates. P.M.R., E.O., and S.C.R. recovered the field *E. hirae* isolate and performed MIC assays with this isolate. All authors reviewed, edited, and approved the final manuscript.

## Acknowledgements

The authors thank Tuesday Simmons for critical review of the manuscript and helpful feedback during manuscript preparation.

---

**Supplementary Figure 1.**
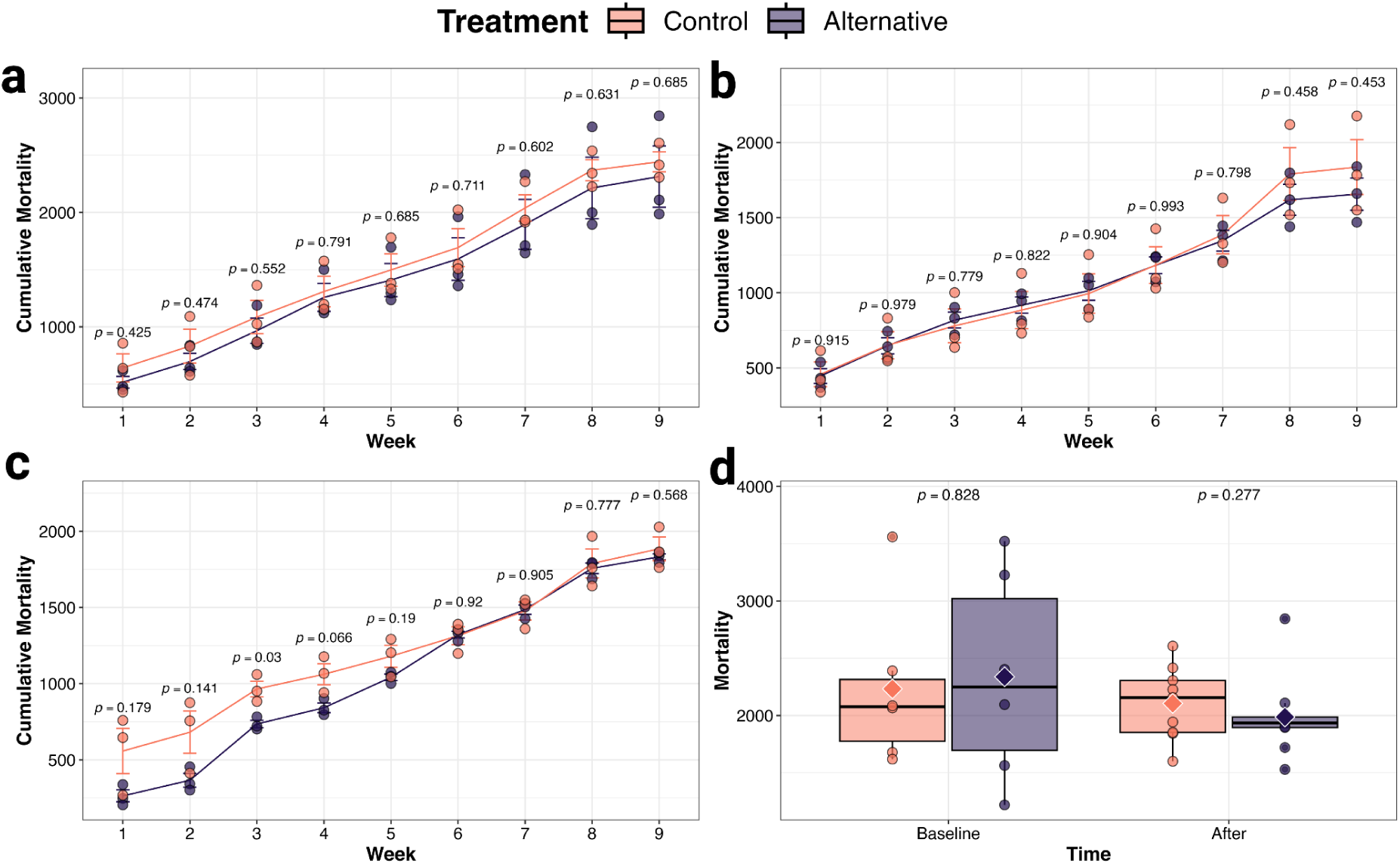
Cumulative mortality estimates across five consecutive growouts. **(a**-**c)** Per-week cumulative mortality for three growouts succeeding study initiation. Points/bars represent the mean ± SEM for each treatment at each week. Within each growout, treatment differences at each week were evaluated using two-sided Welch’s t-tests. **(d)** Total cumulative mortality at week 9 for growouts conducted before (“Baseline”) and after (“After”) study initiation (*n* = 12 and 18, respectively). Treatment effects were evaluated using two-sided Welch’s t-tests for baseline growouts and a generalized linear mixed model (GLMM) for post-initiation growouts, with treatment as a fixed effect and growout as a random intercept. GLMM assumptions and fit were evaluated using the diagnostic procedures described in the main text.

**Supplementary Figure 2.**
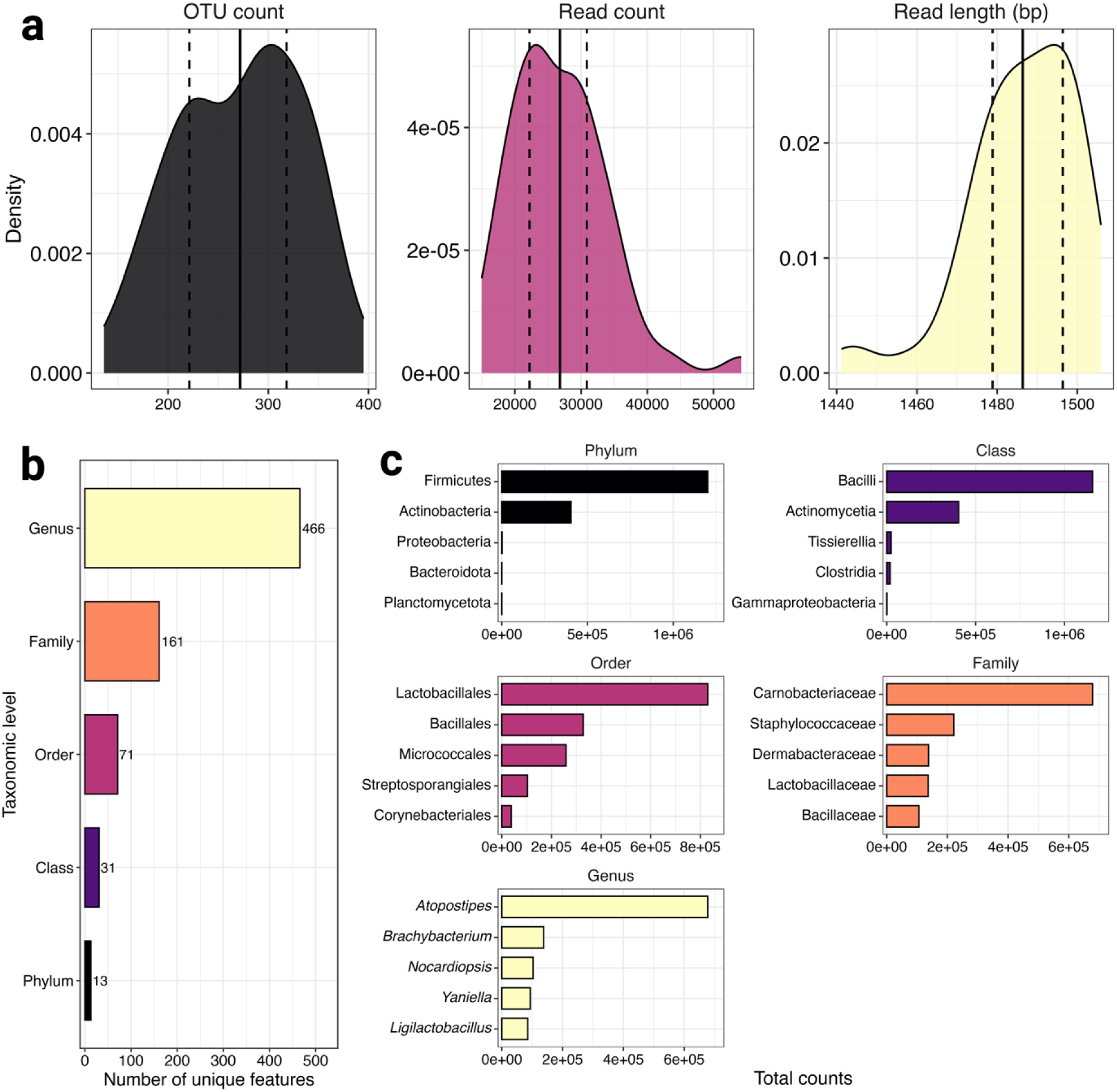
Overview of the 16S rRNA gene sequencing dataset. **(a)** Distribution of per-sample feature sequence count, per-sample read count, and read length. Dashed lines reflect first and third quartiles, and the solid line reflects the mean. **(b)** Number of unique features at phylum-genus taxonomic levels. **(c)** The five most abundant features for each taxonomic level based on counts.

**Supplementary Figure 3.**
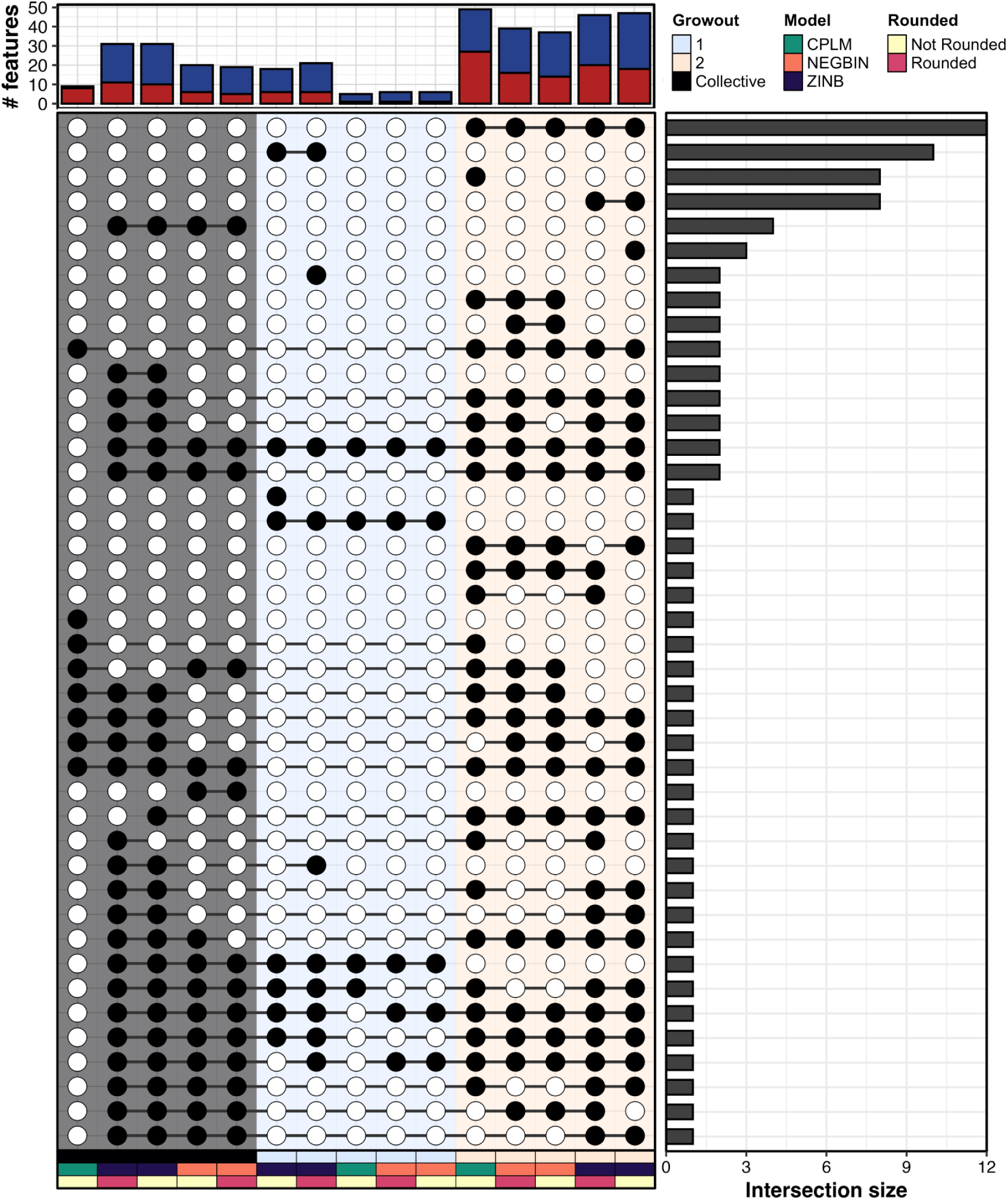
Upset plot of differentially abundant features. The plot shows frequencies and intersections of differentially abundant features (*q* ≤ 0.05) detected across differential abundance models. Column annotations indicate growout, statistical model, and data rounding.

**Supplementary Table1.**
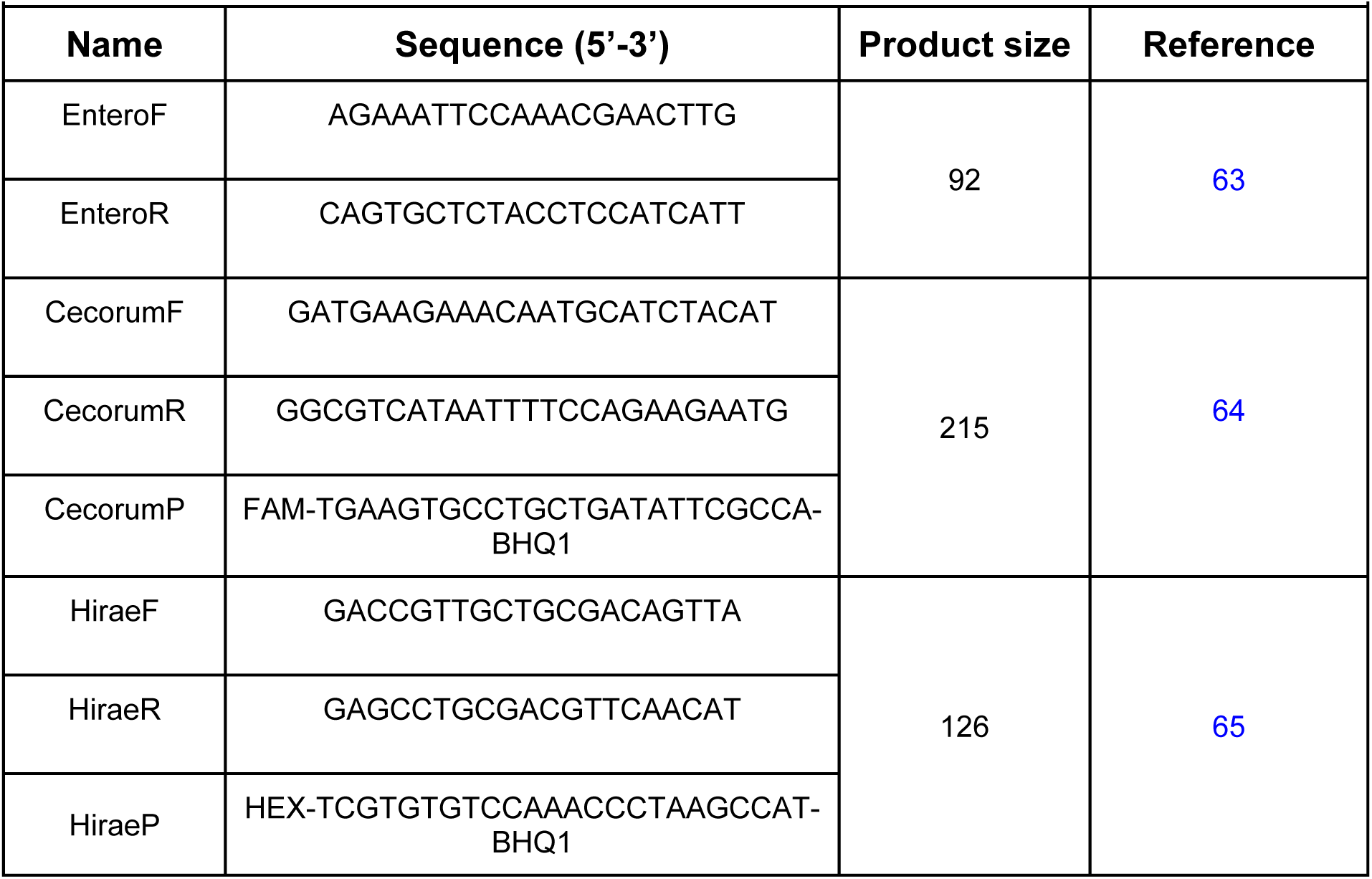
Primer and probe sequences for *Enterococcus* dPCR assays.

